# A lysosomal dimmer switch regulates cellular quiescence depth

**DOI:** 10.1101/413989

**Authors:** Kotaro Fujimaki, Ruoyan Li, Hengyu Chen, Kimiko Della Croce, Hao Helen Zhang, Jianhua Xing, Fan Bai, Guang Yao

## Abstract

Numerous physiological and pathological phenomena are associated with the quiescent state of a cell. Cellular quiescence is a heterogeneous resting state; cells in deep than shallow quiescence require stronger growth stimulation to exit quiescence and reenter the cell cycle. Despite the importance of quiescent cells such as stem and progenitor cells to tissue homeostasis and repair, cellular mechanisms controlling the depth of cellular quiescence are poorly understood. Here we began by analyzing transcriptome changes as rat embryonic fibroblasts moved progressively deeper into quiescence under increasingly longer periods of serum starvation. We found that lysosomal gene expression was significantly upregulated in deep than shallow quiescence, which compensated for gradually reduced autophagy flux observed during quiescence deepening. Consistently, we show that inhibiting lysosomal function drove cells deeper into quiescence and eventually into a senescence-like irreversibly arrested state. By contrast, increasing lysosomal function progressively pushed cells into shallower quiescence. That is, lysosomal function modulates quiescence depth continuously like a dimmer switch. Mechanistically, we show that lysosomal function prevents quiescence deepening by reducing oxidative stress in the cell. Lastly, we show that a gene expression signature developed by comparing deep and shallow quiescent cells can correctly classify senescent and aging cells in a wide array of cell lines *in vitro* and tissues *in vivo*, suggesting that quiescence deepening, senescence, and aging may share common regulatory mechanisms.

## INTRODUCTION

A salient characteristic of multicellular organisms is the tight regulation of cell division. Cells in the body proliferate upon proper growth signals and remain dormant in the absence of such signals. The dormant state, reversible to proliferation, is referred to as cellular quiescence, which is fundamental to many physiological phenomena such as stem cell homeostasis and tissue repair^1-4^. Consequently, dysregulation of cellular quiescence can lead to a range of hyper- and hypo-proliferative diseases including cancer and aging^5-9^.

Quiescent cells can progress into deeper quiescence, from which they require stronger growth stimulation and a longer time to reenter the cell cycle. Deep quiescence, after all, can still revert to proliferation, making it phenotypically distinct from other irreversibly arrested cellular states such as senescence (Fig. S1a). *In vitro*, deep quiescence arises when cells are cultured longer under quiescence inducing signals such as contact inhibition^10, 11^ and serum starvation^12^. *In vivo*, deep quiescence is associated with aging—e.g., hepatocytes in older than younger rats take a longer time to reenter the cell cycle upon partial hepatectomy^13, 14^. On the other hand, cells can shift into shallower quiescence and become sensitized to growth stimulation, as seen in neural stem cells and muscle satellite cells post injury^15, 16^. Despite the importance of quiescence depth for tissue repair and regeneration, cellular mechanisms regulating quiescence depth are poorly understood.

In this study, we set out to investigate what regulates quiescence depth in a rat embryonic fibroblast (REF) cell model. We identified sequential transcriptome changes as cells moved progressively deeper into quiescence under longer-term serum starvation. In particular, we found that lysosomal gene expression and biogenesis continuously increased with quiescence deepening; autophagy flux, however, decreased.

Lysosome, the hydrolytic enzyme-filled organelle in the cell, functions to break down many types of biomolecules including proteins, nucleic acids, carbohydrates, and lipids through processes such as autophagy and endocytosis. Lysosomal function has been shown to prevent irreversible cellular states such as senescence, terminal differentiation, and apoptosis^6, 17-20^. Here we found that increased lysosomal gene expression and expanded lysosomal biogenesis in deep quiescent cells were compensatory for decreased lysosomal function. We show that lysosomal function, like a dimmer switch, continuously regulates quiescence depth and thus the proliferative potential of quiescence cells, by reducing the accumulation of intracellular reactive oxygen species (ROS). Furthermore, we found that a gene expression signature developed by comparing deep and shallow quiescent REF cells was able to correctly classify senescent and aging cells in a wide array of cell lines *in vitro*^21, 22^ and tissues *in vivo*^23, 24^, suggesting the existence of shared regulatory mechanisms underlying these cell fates and a possible sequential transition from shallow to deep quiescence and eventually to irreversible senescence that may contribute to aging.

## RESULTS

### Transcriptome changes during quiescence deepening

Similar to our previous observation^12^, REF cells moved into deeper quiescence progressively with longer-term serum starvation. After a 2-day serum starvation, the entire cell population entered quiescence as demonstrated by their negative DNA incorporation of 5-ethynyl-2’-deoxyuridine (EdU) and a complete shut- off of E2f1expression (Fig. S1b). E2f1 is a member of the E2f family of transcription factors; it up-regulates a large battery of genes involved in DNA replication and cell cycle progression, and acts as an effector of an Rb-E2f bistable switch that controls the all-or-none transition from quiescence to proliferation^25, 26^. With increasingly longer serum starvation, cells moved deeper into quiescence, as shown by their longer average “waiting time” before resuming DNA replication upon serum stimulation. For example, it took 19, 22, and 25 hours of serum stimulation for ∼60% of the cells that were under 4-, 8-, and 14-day serum starvation (labelled as STA in figures for simplicity), respectively, to become EdU^+^ (Fig. 1a); after 19 hours of serum stimulation, while ∼80% of 2-day serum-starved cells became EdU^+^, only ∼10% of 14-day serum-starved cells did so (Fig. 1a). Importantly, deep quiescent cells were not irreversibly arrested; upon serum stimulation, they were able to re-enter the cell cycle (EdU^+^ and E2f-ON, Fig. S1b).

**Figure 1.**
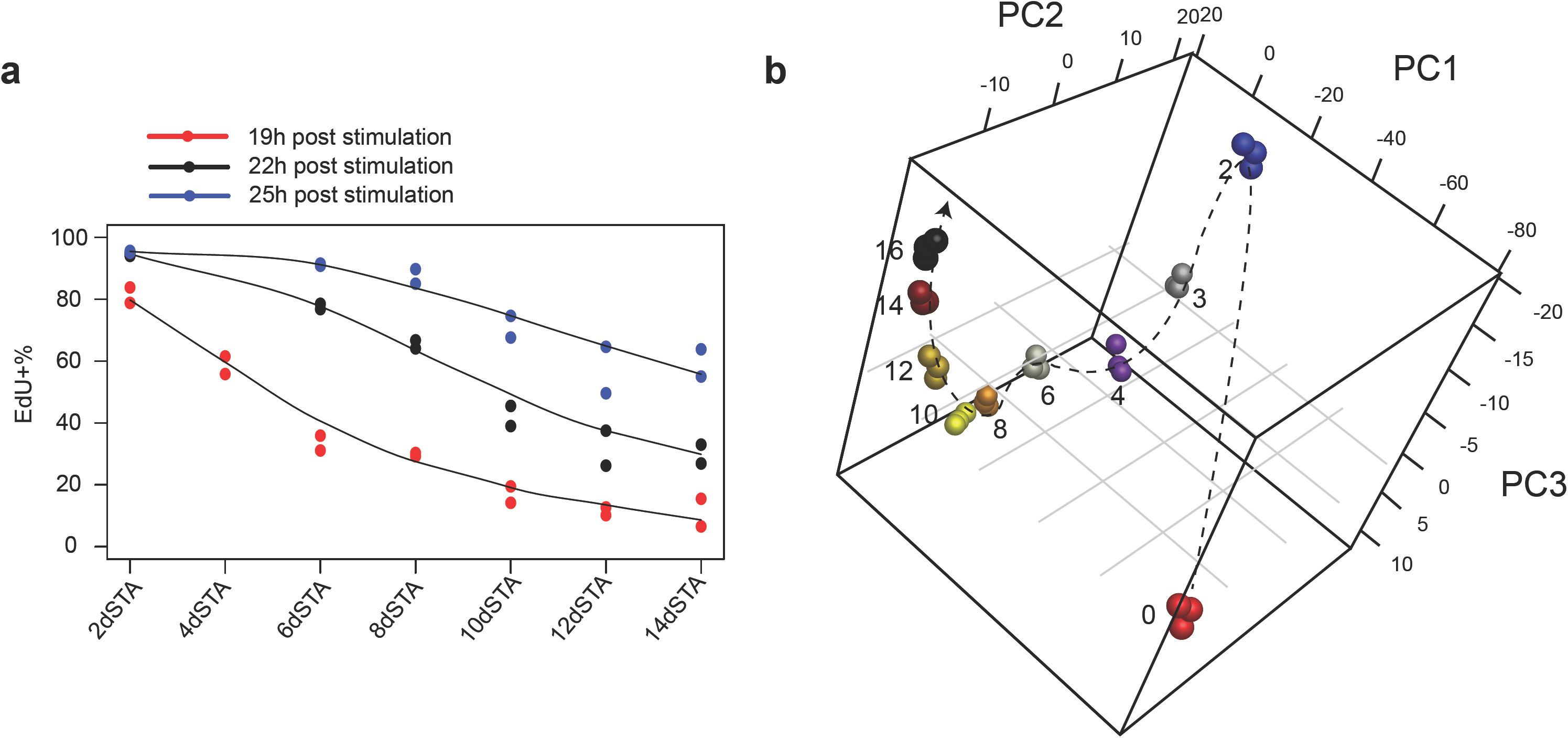
Sequential quiescence deepening and transcriptome change in serum-starved REF cells over time. (**a**) EdU^+^% of 2- to 14-day serum-starved cells upon serum stimulation. Cells in duplicated wells (n = 2) were harvested 19, 22, and 25 hours after stimulation with serum (20%) and subjected to EdU assay. (**b**) Principal component analysis of proliferating and 2- to 16-day serum-starved cells was performed based on 2,736 differentially expressed genes in RNA-seq time course. Days of serum starvation are indicated next to the sample (triplicates per condition). Dashed arrow is plotted for clarity.

To better understand molecular mechanisms regulating quiescence depth at the transcriptional level, we performed RNA-seq analysis of 0- to 16-day serum-starved cells. As expected, quick down-regulation (within 2-day serum starvation) was observed in the expression of well-characterized proliferation genes such as E2f1, Cdk2, and Cdk4 (Fig. S2a), E2f1 target genes (Fig. S2b), and proliferation related gene clusters (#7 and 9, Fig. S2c). Conversely, the expression of growth inhibitory genes, such as Rb1, Cdkn1a (p21^Cip1^), and Cdkn1b (p27^Kip1^), was up-regulated upon serum starvation (Fig. S2a). The expression of Cdkn2a (p16^INK4A^), a senescence marker^27^, remained low as cells moved deeper in quiescence (3- to 16-day serum starvation, Fig. S2d), consistent with quiescence being a reversible state.

In terms of the global gene expression profile, it not only changed drastically when cells transitioned from proliferation to quiescence (0- to 2-day serum starvation) but kept changing sequentially as cells moved from shallow to deep quiescence (2- to 16-day serum starvation) (Fig. 1b). This sequential change was reflected in 9 gene clusters that exhibited different temporal dynamic patterns (Fig. 2a, b). In particular, the expression of one gene cluster increased progressively without plateauing as cells moved deeper into quiescence (cluster 1, Fig. 2b). In this quiescence deepening-associated gene cluster, multiple biological functions were enriched; among them, the enrichment of lysosomal genes was the most statistically significant (Fig. 2c and S2c). Consistently, lysosomal genes had the strongest positive correlation with deep quiescence in Gene Set Enrichment Analysis (GSEA, NES = 3.12; Fig. 2d).

**Figure 2.**
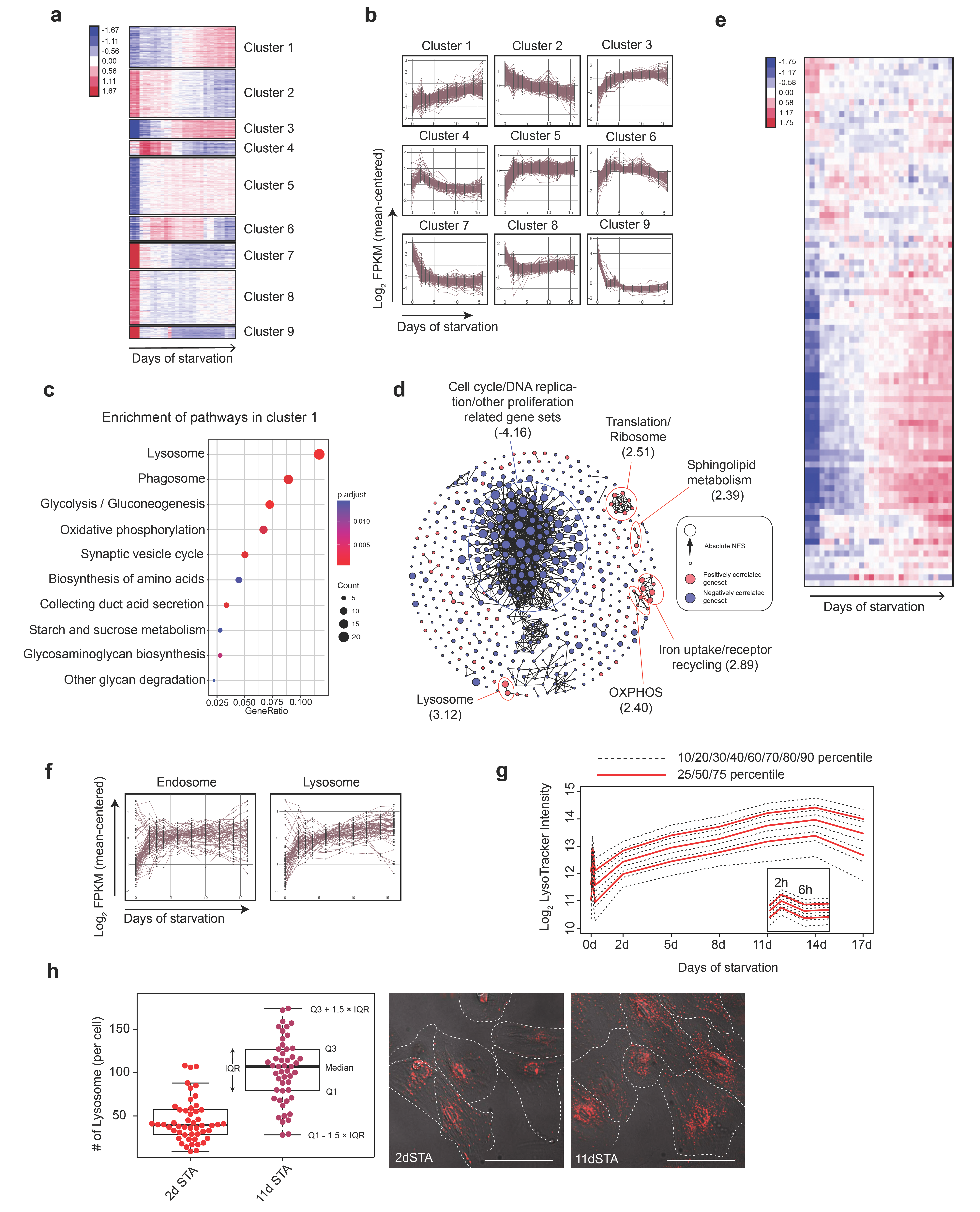
Lysosomal gene expression and biogenesis increase along quiescence deepening. K-means clustering of 2,736 differentially expressed genes in RNA-Seq analysis (0- to 16-day serum-starved cells). Sample columns (0, 2, 3, 4, 6, 8, 10, 12, 14, and 16-day serum starvation) are ordered chronologically with three replicates at each time point. Up- and down-regulation of genes are shown in red and blue, respectively; color gradient bar indicates the degree of change in gene expression (normalized and log transformed, see Methods for details). Gene expression dynamics within each K-means cluster in **a**. Expression of each gene in a given cluster is shown in a time-course curve. (**c**) Pathways enriched (p < 0.05) in cluster 1. GeneRatio (x-axis) and dot size indicate the fraction and number of genes involved in the indicated pathway in cluster 1, respectively. Color gradient bar indicates statistical significance of pathway enrichment in KEGG over-representation test based on p-values adjusted by Benjamini correction. (**d**) Gene sets significantly correlated to quiescence depth (FDR < 0.1) in GSEA analysis. Gene sets are connected by edges based on their similarity (Jaccard index ≥ 0.5) when applicable. Gene sets positively and negatively correlated to quiescence deepening are shown in red and blue nodes, respectively. Node size represents the absolute Normalized Enrichment Score (NES) in the GSEA result. Biological functions that are the most negatively and positively (top 5 in statistical significance) correlated with quiescence deepening are demarcated by blue and red circles, respectively; the maximum NES value of all enclosed nodes in a circle is shown in parenthesis. (**e**) Heat map of time-course expression of lysosomal genes in RNA-Seq analysis (0- to 16-day serum-starved cells). Sample columns are ordered chronologically (as in **a)** and rows are ordered by hierarchical clustering. Color gradient bar indicates the degree of change in gene expression as in **a**. (**f**) Time-course expression of endosomal and lysosomal differentially expressed genes in RNA-Seq analysis (0- to 16-day serum-starved cells). (**g**) LysoTracker intensity distribution in REF cells serum starved from 0 to 17 days (0 h, 2 h, 6 h, 2 d, 5 d, 8 d, 11 d, 14 d, and 17 d). Cellular LysoTracker intensity measured by flow cytometry was log_2_ transformed and plotted with the indicated percentiles (agglomeration of three replicates, each with over 10,000 cells at a time point). (**h**) LysoTracker foci number per cell in 2- and 11-day serum-starved cells (n = 50 each). Box plot: Q1, Q3 refer to the 1^st^ and 3^rd^ quantiles, respectively; IQR, interquartile range = Q3 - Q1; the same below unless otherwise noted. (Inset) Representative lysoTracker foci microscopy. Each red dot represents a LysoTracker-stained lysosome in the cell. Scale bar = 50 µm.

### Lysosomal gene expression and biogenesis increase as quiescence deepens

The majority of lysosomal genes, encoding for various lysosomal enzymes, activator proteins, membrane proteins, and ion channel proteins, increased their expression as cells moved into deeper quiescence (Fig. 2e and Fig. S3). In comparison, the expression of most genes associated with endosome, another cellular organelle in the endosomal-lysosomal system, did not increase significantly as quiescence deepened (Fig. 2f).

Lysosomal biogenesis also increased in deep quiescence. Lysosomal mass first showed a brief pulsatile adaptive response (within the first 6 hours) upon serum starvation and then continuously increased during the following 14 days, as seen from the stained LysoTracker intensity (Fig. 2g). The initial pulsatile response of lysosomal mass was likely due to an adaptive mTOR-autophagy response to serum starvation as previously reported^28^. The continuously increased lysosomal mass in deep quiescence was at least partially due to increased lysosomal number; e.g., 11-day serum-starved cells exhibited significantly more lysosomal foci than 2-day serum-starved cells (> 2.5-fold, p = 4.9*e^-14^ in a one-tailed *t*-test; Fig. 2h).

### Inhibiting lysosomal function deepens cellular quiescence

Were increased lysosomal gene expression and biogenesis responsible for driving cells into deeper quiescence, or were they a consequence of such deepening? If increased lysosomal gene expression and biogenesis drove cells into deeper quiescence, inhibiting lysosomal function would prevent such deepening (*I*, Fig. S4a); if they were merely downstream effects caused by quiescence deepening, inhibiting lysosomal function may not affect quiescence depth (*II*, Fig. S4a); however, if they were downstream effects that were compensatory for and preventing further quiescence deepening, inhibiting lysosomal function would lead to even deeper quiescence (*III*, Fig. S4a).

In order to test these competing hypotheses, we first performed pharmacological inhibition of lysosomal function and measured corresponding quiescence-depth change (Fig. 3a). We used two lysosomal inhibitors, bafilomycin A1 (Baf) and chloroquine (CQ) that prevent lysosomal acidification^29^, and found that both drugs inhibited lysosomal function as expected, evident by impaired proteolytic degradation within the lysosomal compartment (Fig. S4b). When quiescent cells were treated with these two lysosomal inhibitors, higher serum concentrations were required to activate E2f1 and initiate DNA replication in a drug dose-dependent manner (Fig. 3b, c; red arrow pointed: serum concentration for activating ∼50% of cells). This lysosomal inhibitor-caused quiescence deepening occurred regardless of the preceding quiescence depth before drug treatment; as shown in Fig. S4c, a higher serum concentration was required to activate drug-treated (blue curve) than non-treated (red curve) cells at all test conditions (serum-starvation days). Furthermore, with CQ treatment at a high concentration (20 μM), the majority of cells entered a senescence-like, irreversibly arrested state and did not re-enter the cell cycle even with strong growth stimulation (20% serum, Fig. 3b, c). This finding that inhibiting lysosomal function deepened quiescence suggests that increased lysosomal gene expression and biogenesis played a compensatory role to prevent further deepening of quiescence.

**Figure 3.**
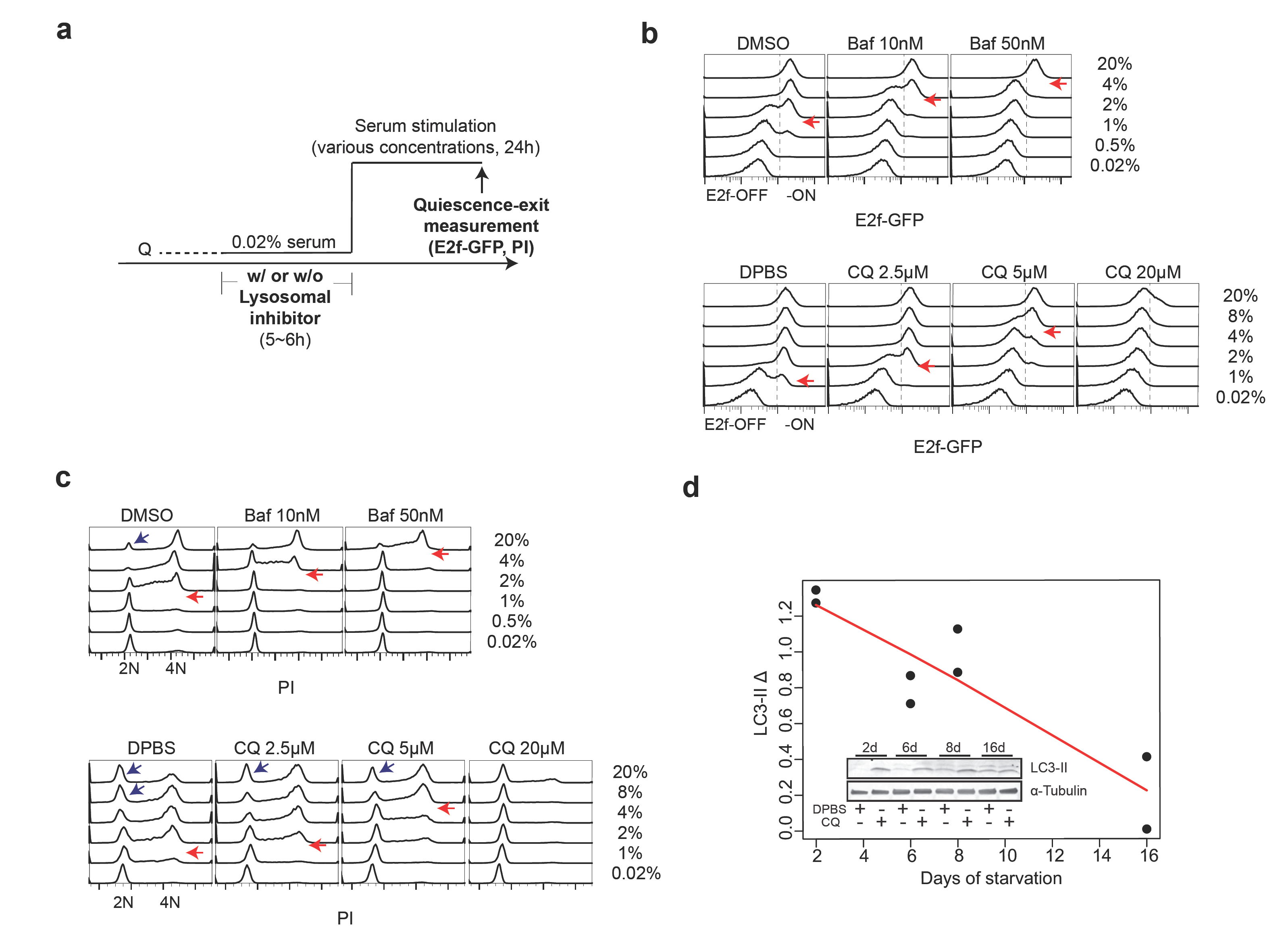
Lysosomal inhibition pushes cells into deeper quiescence as autophagy declines in longer-term serum-starved cells. (**a**) Experimental scheme of quiescence-depth assay with lysosomal inhibitor treatment. Q, quiescent state. (**b, c**) Readouts of E2f-GFP (**b**) and PI staining in quiescence-depth assay performed on 2-day serum-starved cells (∼10,000 cells per sample, with the highest frequency set to 100% at the y-axis of each histogram). Cells at E2f-ON and - OFF states (**b**) and G0/G1 (2N), S (2-4N), and G2/M phase (4N) (**c**) are as indicated. Red arrows indicate serum concentrations that activated E2f1 or initiated DNA replication in approximately half of the cell population. Note that each blue arrow-pointed peak in **c** contained a subset of recently divided cells following strong serum stimulation. (**d**) LC3-II turnover assay in 2, 6, 8, 16-day serum-starved cells. Quantified LC3-II Δ (difference between LC3-II signal intensity normalized against α-Tubulin control in CQ-treated and non-treated cells) in duplicate samples is shown with a linear fit (red line). (Inset) A representative immunoblot image.

We hypothesized that increased lysosomal gene expression and biogenesis in deep quiescence might respond to and compensate for a decreased lysosomal function. Indeed, we found that autophagy flux, an indicator for a primary lysosomal function, gradually declined as cells moved deeper into quiescence as seen in LC3-II turnover assay^30^ (Fig. 3d). We reasoned that a decreased lysosomal function in longer-term serum-starved cells, which was partially but not completely compensated by increased lysosomal gene expression and biogenesis, was responsible for quiescence deepening.

### Enhancing lysosomal function pushes cells toward shallower quiescence

If a decreased lysosomal function is responsible for quiescence deepening, enhancing such function would likely counteract this trend and push cells into shallower quiescence. Consistent with this hypothesis, we observed that nutrient starvation, a known inducer of autophagy, pushed REF cells into shallower quiescence (higher E2f-ON% upon serum stimulation in cells treated with PBS over control, Fig. S5a), which is similar to a previous observation in quiescent NSCs^31^. However, nutrient starvation induces a range of cellular responses (e.g., AMPK activation and mTOR inhibition) in addition to autophagy. To increase lysosomal function directly, we considered to enhance lysosomal gene expression and biogenesis in REF cells. Lysosomal gene expression and biogenesis can be up-regulated by an MiT/TFE family of transcription factors Tfeb, Mitf, and Tfe3, which bind to a CLEAR-box sequence upstream of many lysosomal genes and are known as the master regulator of lysosomal biogenesis^32-35^. Remarkably, the expression of Mitf and Tfe3 but not Tfeb increased significantly in deep quiescence (Fig. 4a; *p* = 0.028, 0.032, and 0.827 for Mitf, Tfe3, and Tfeb, respectively, in a one-tailed *t*-test comparing 16-day and 2-day serum-starved cells); meanwhile, Mitf but not Tfeb and Tfe3 showed a high degree of co-expression with lysosomal genes in quiescence (Fig. S5b). Together, these results suggested a unique role for Mitf in regulating lysosomal function in the quiescent REF cell model. Indeed, we found that ectopic Mitf expression in quiescent REF cells enhanced lysosomal biogenesis (higher LysoTracker intensity, Fig. 4b, c) and autophagy flux (higher LC3-II Δ, Fig. 4d, e), and meanwhile pushed cells into shallower quiescence (higher EdU+% upon serum stimulation, Fig. 4f). Notably, there was a monotonic correlation between the level of introduced Mitf expression vector (indicated by mCherry intensity, x-axis, Fig. 4g) and the EdU+ level upon serum stimulation (normalized to mCherry control, z-axis, Fig. 4g), suggesting that cells can be continuously driven to shallower quiescence by enhancing lysosomal function.

**Figure 4.**
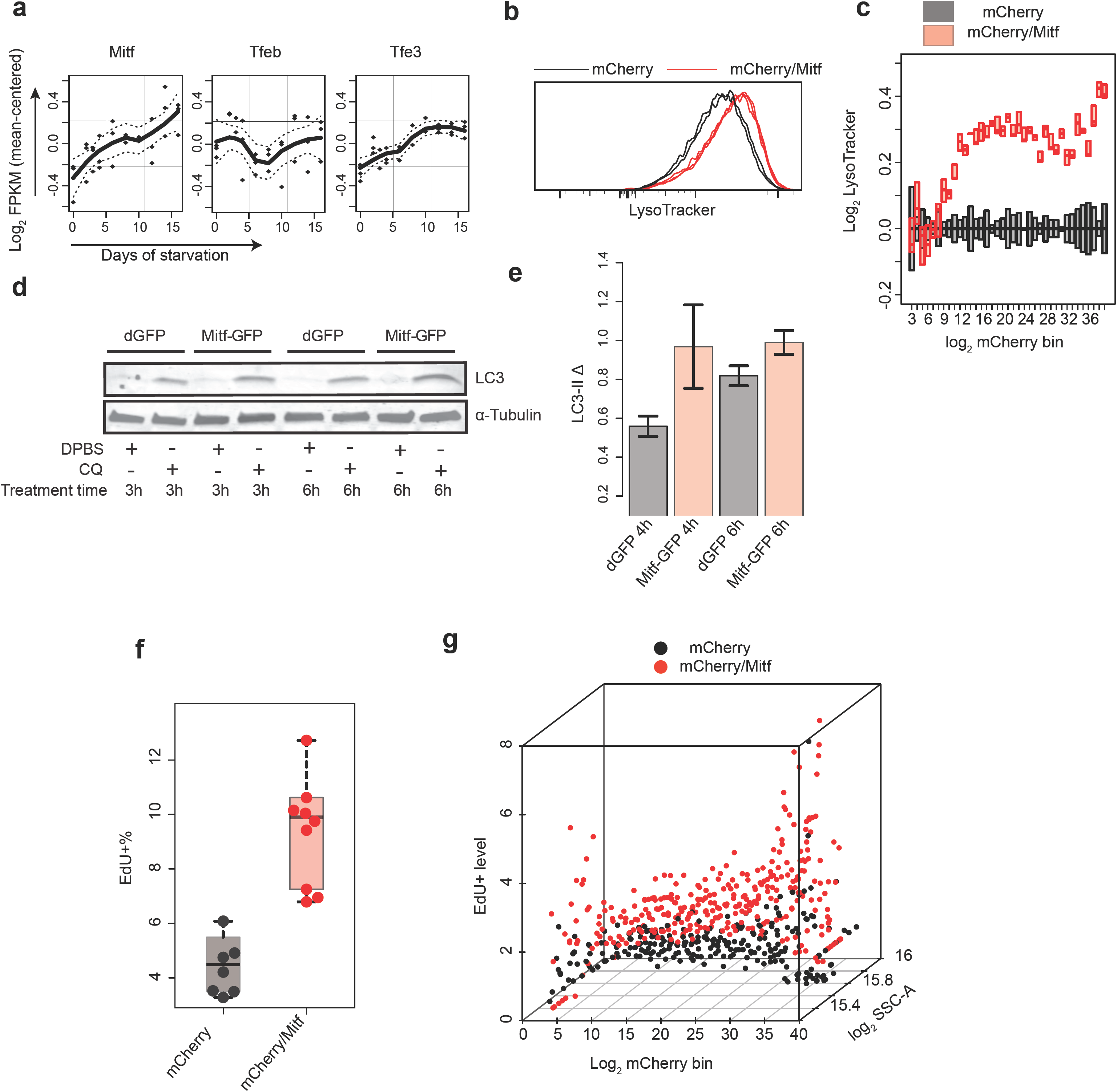
Enhancing lysosomal function pushes cells toward shallower quiescence. (**a**) Time-course expression of MiT/TFE family members in RNA-Seq analysis (0- to 16-day serum-starved cells). Dash line, s.e.m of fitted line. (**b, c**) Lysosomal mass indicated by LysoTracker intensity. Cells were co-transfected with Mitf and mCherry expression vectors (mCherry/Mitf, triplicates) or mCherry control alone (mCherry, duplicates) and induced to quiescence by 4-day serum starvation. (**b**) Distribution of LysoTracker intensity in ∼10,000 cells per sample, with the highest frequency set at 100% (y-axis). (**c**) Binned LysoTracker intensity (log transformed with the median of mCherry control set at 0; bottom and top edges of each box indicate the 1^st^ and 3^rd^ quartiles, respectively). Cells were grouped according to their mCherry intensity (log transformed) into 40 even-width bins (bins with cell number < 10 were filtered out). The mCherry intensity indicates co-transfected Mitf vector level^12^ in mCherry/Mitf cells. (**d, e**) LC3-II turnover assay. Cells were transfected with Mitf expression vector or dGFP control and induced to quiescence by 4-day serum starvation. (**d**) Representative immunoblot image. (**e**) Quantified LC3-II Δ (between CQ-treated and non-treated cells as in Fig. 3d, duplicates). Error bar, s.e.m. (**f, g**) Quiescence-depth assay in mCherry or mCherry/Mitf transfected cells. Transfected cells were induced to quiescence by 4-day serum starvation and stimulated with 0.5% serum. **(f)** EdU^+^% measured 24 hours after the initiation of serum stimulation (in 7 and 9 mCherry and mCherry/Mitf transfection, respectively, ∼10,000 cells each). (**g**) Transfected cells in **f** were binned according to mCherry intensity as in **c**. EdU^+^% of mCherry/Mitf cells was normalized to that of mCherry control cells in each bin. The resultant “EdU+ level” (y-axis) indicates relative fold increase of quiescence exit (EdU^+^) associated with ectopic Mitf level (indicated by mCherry intensity, x-axis) over corresponding mCherry control. Each dot in a given bin corresponds to a transfected cell population in **f**.

### Lysosomal function prevents quiescence deepening via ROS reduction

Lysosomes are known to play roles in antioxidation and energy generation in quiescent stem cells^6, 17, 18, 20^. Thus, here we tested whether lysosomal function potentially prevents quiescence deepening by either of the two mechanisms. If it does, we reasoned, increasing antioxidation and/or energy generation in the cell would reduce quiescence depth. To this end, we first supplemented serum-starved quiescent cells with cell-permeable methyl pyruvate (MPy) that enhances energy generation, or with antioxidant 2-mercaptoethanol (βME), to test whether cells may move to shallower quiescence in response. We found that supplementing MPy to 2-day serum-starved cells for additional 2-4 days did not reduce quiescence depth with statistical significance (Fig. 5a and Fig. S6a) and even increased quiescence depth at high MPy concentrations (3.6 and 10 mM, Fig. S6b), indicating that energy generation may not be responsible for how lysosomal function prevents quiescence deepening in the tested time window.

**Figure 5.**
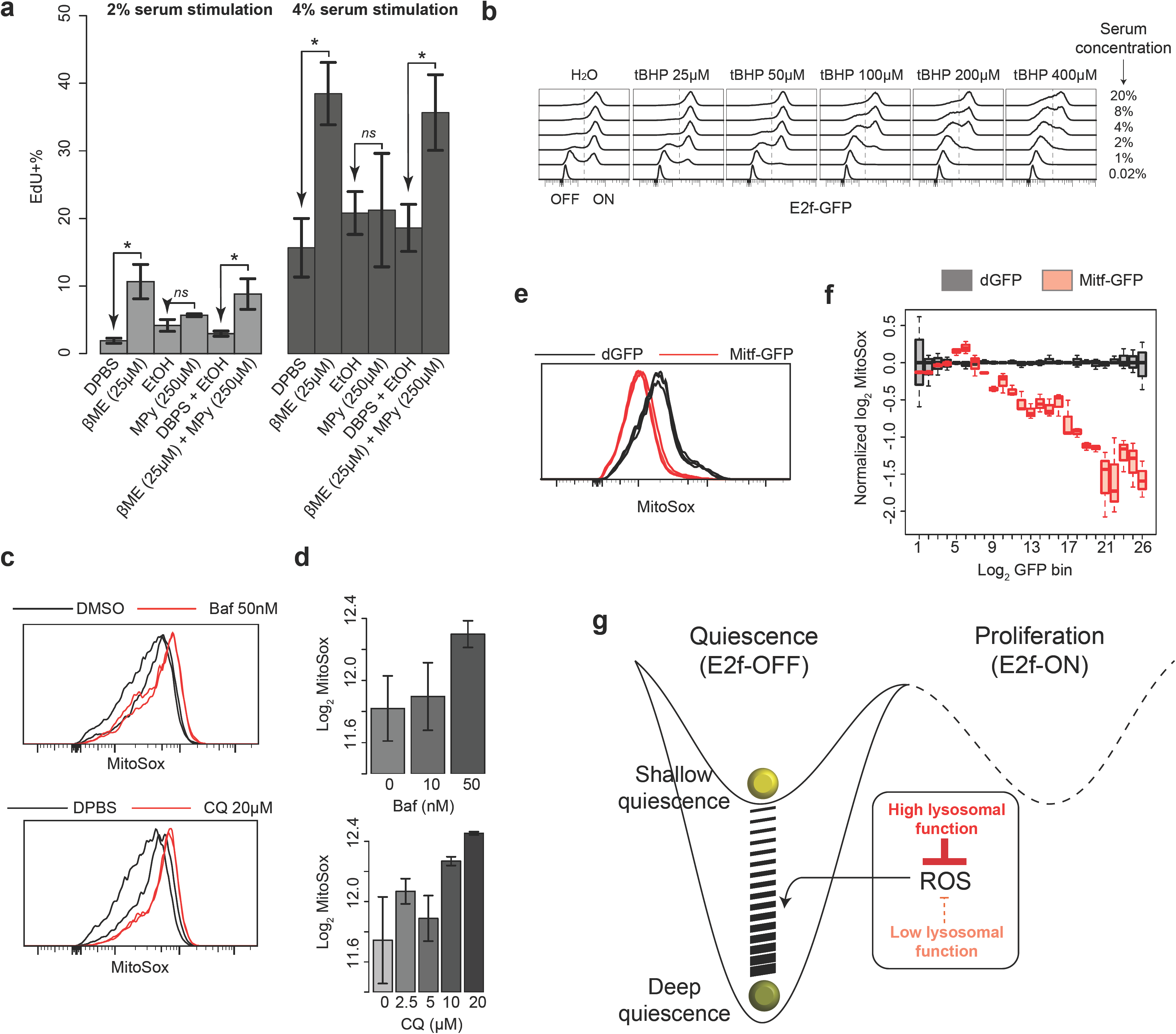
Lysosomal function reduces oxidative stress and prevents quiescence deepening. (**a**) 2-day serum-starved cells were further cultured in starvation medium containing βME and/or MPy for 4 days and stimulated with 2% or 4% serum for 24 hours, followed by EdU assay. Star sign (*) and *ns*, statistical significance (p < 0.05) and insignificance, respectively, in a one-tailed (arrow pointed) *t*-test comparing the average EdU^+^% of βME- and/or MPy-treated versus non-treated cells (triplicates with ∼10,000 cells each at either condition). Error bar, s.e.m. (**b)** 2-day serum-starved cells were treated with tBHP at indicated concentrations for 1 hour and stimulated with various concentrations of serum as indicated for 24 hours, followed by E2f-GFP assay. (**c,d**) Intracellular ROS level in quiescent REF cells treated with lysosomal inhibitors. 2-day serum-starved cells were treated with Baf or CQ at indicated concentrations for 6 hours and subjected to MitoSox measurement (duplicates with ∼10,000 cells each). MitoSox signal distribution with the highest frequency set at 100% (y-axis, **c)** and its median (log transformed, **d**) are shown. Error bar, s.e.m. (**e, f**) Intracellular ROS level in quiescent REF cells transfected with Mitf expression vector. Cells were transfected with Mitf-GFP or dGFP control vectors and induced to quiescence by 4-day serum starvation, followed by MitoSox measurement (triplicates with ∼10,000 cells each). (**e)** MitoSox signal distribution with the highest frequency set at 100% (y-axis). (**f**) Binned MitoSox intensity (log transformed and normalized with the median of dGFP control set at 0). Cells were grouped according to their log transformed GFP intensity (positively correlated with Mitf level) into 26 even-width bins. (**g**) Model of quiescence-depth regulation by lysosome function.

In comparison, antioxidation appears to play an important role in the lysosomal regulation of quiescence depth. First, supplementing antioxidant βME drove quiescent cells into a shallower state, from which cells became more sensitive to growth signals (higher EdU+% upon serum stimulation, Fig. 5a); conversely, supplementing an oxidative reagent tert-butyl hydroperoxide (tBHP) drove cells into deeper quiescence (lower E2f-ON% upon stimulation, e.g., at 2% serum, with increasing tBHP concentration, Fig. 5b). These results indicate that the degree of oxidative stress is positively correlated with quiescence depth. Next, we found that inhibiting lysosomal function by Baf and CQ, which deepened quiescence, increased mitochondrial ROS level (higher MitoSox intensity, Fig. 5c, d). Conversely, enhancing lysosomal function by ectopic Mitf expression, which reduced quiescence depth, suppressed mitochondrial ROS (lower MitoSox intensity, Fig. 5e) in a dose-dependent manner (Fig. 5f). Together, our findings suggest that a) lysosomal function reduces intracellular ROS, the mediator of oxidative stress; b) ROS reduction helps prevent quiescence deepening; and c) the change of lysosomal function continuously modulates quiescence depth, acting like a dimmer switch (Fig. 5g).

### Quiescence deepening parallels cellular senescence and aging

Lastly, we examined whether and how deep quiescence in REF cells is related to the quiescent states of other cell types (case A) and other non-growth cellular states such as senescence (case B). To this end, we first developed a gene expression signature indicating the quiescence depth in REF cells by performing a linear regression analysis of the time-course RNA-seq data (2-day to 16-day serum starvation, shallow to deep quiescence). We next applied this gene signature to other publicly available RNA-seq datasets related to cases A and B above.

As an example of case A, dormant neural stem cells (NSCs) *in vivo* transition into a shallower quiescent state called primed NSCs upon neural injury^16^. When we applied the quiescence-depth gene signature of REF cells to the RNA-seq data corresponding to NSCs before and after neural injury, we obtained quiescence depth scores (QDS) from the linear regression model to indicate the relative quiescence depth of NSCs. We found that the QDS of primed NSCs after injury was significantly smaller (i.e., shallower quiescence) than that of dormant NSCs before injury (Fig. 6a). This result suggests that the quiescence-depth signature of REF cells *in vitro* can predict the relative quiescence depth of NSCs *in vivo*, and that this gene signature may reflect shared quiescence regulatory mechanisms across different cell types in different microenvironments.

**Figure 6.**
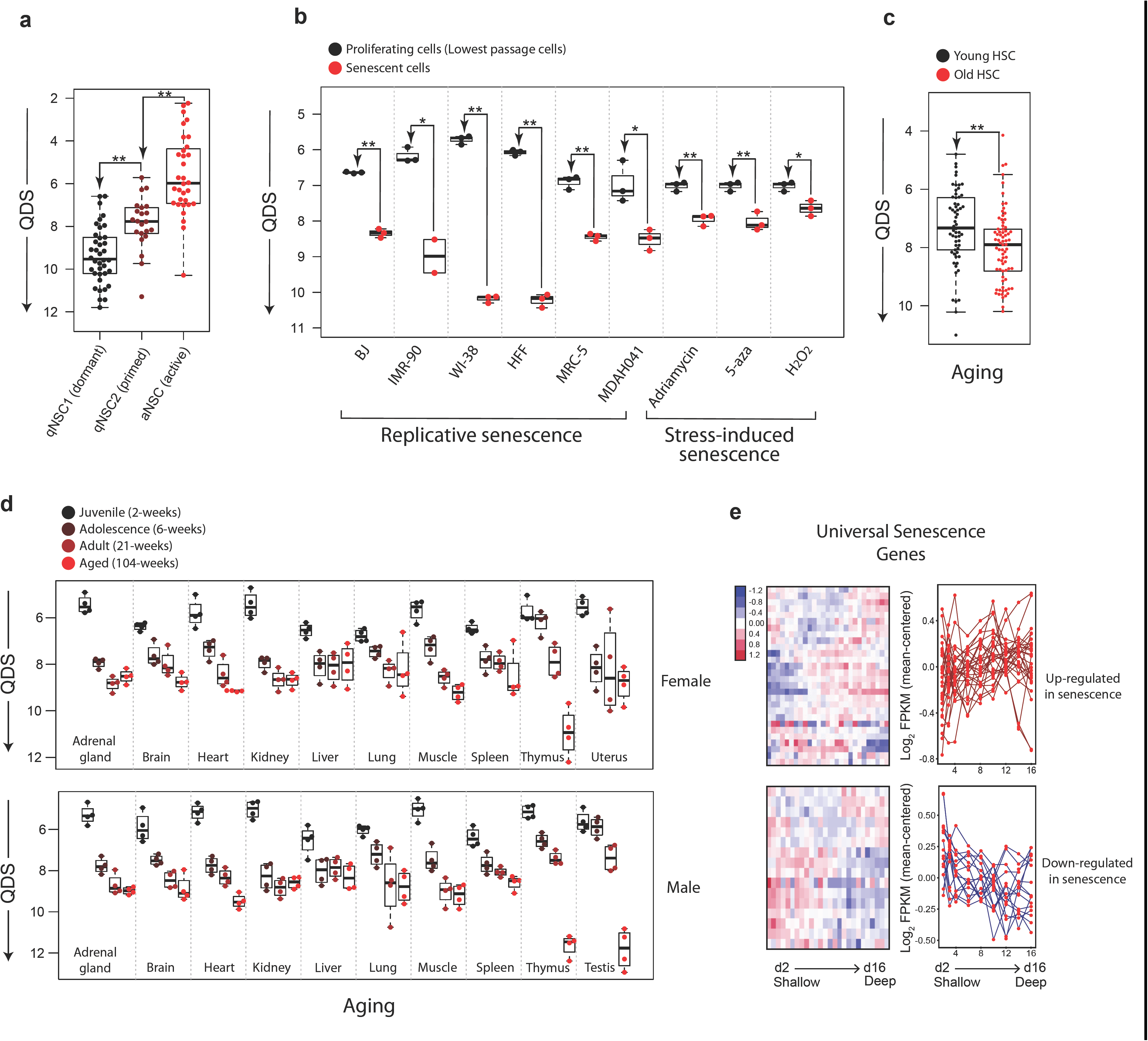
Quiescence deepening parallels cellular senescence *in vitro* and aging *in vivo*. (**a)** QDS of dormant, primed, and activated NSCs predicated based on the quiescence-depth gene expression signature developed in REF cells. See Methods for details. * and ** indicate p-value < 0.05 and 0.01 in a one-tailed *t*-test, respectively. (**b)** Predicted QDS of indicated cell lines at proliferative state versus replicative senescence or stress-induced senescence (caused by DNA damaging agent Adriamycin, DNA demethylation agent 5-aza-2-deoxycytidine (5-aza), or oxidant H2O2). (**c**) Predicted QDS of hematopoietic stem cells (HSCs) collected from young (2-3 months) or aged (20-25 months) mice. (**d**) Predicted QDS of indicated organ types harvested from rats (female, top; male, bottom) at indicated age stages. (**e**) Expression dynamics of universal senescence genes in RNA-Seq analysis (2-to 16-day serum-starved cells), shown in a hierarchical clustering heatmap (left) or an overlaid line plot (right). Universal senescence genes up- or down-regulated in senescent cells are grouped in top and bottom panels, respectively.

To test case B, we analyzed RNA-seq datasets associated with cellular senescence and aging by applying the QDS model with the quiescence-depth gene signature of REF cells. Similar to deep quiescence, senescence and aging are at least partially driven by ROS and can be counteracted by the lysosome-autophagy pathway^6, 8, 18, 19, 36^. With this mechanistic similarity in mind, it was still striking to us that QDS correctly predicted cellular senescence and aging in a wide array of cell lines *in vitro*^21, 22^ and tissues *in vivo*^23, 24^ (Fig. 6b-d). For example, QDS was significantly larger in the senescent cell populations than in proliferating controls in all six cell lines (BJ, IMR-90, WI-38, HFF, MRC-5, and MDAH041) studied under replicative senescence and with all three inducers (Adriamycin, 5-aza, and H_2_O_2_) studied under stress-induced senescence (Fig. 6b). Consistently, a set of “universal senescence genes” identified from meta-analysis^37^ exhibited similar expression changes (i.e., up- or down-regulation) under both deep quiescence and senescence (Fig. 6e). Furthermore, significantly increased QDS was associated with the aging of hematopoietic stem cells (Fig. 6c) and of all eleven studied rat organs (Fig. 6d). The common increase in QDS observed in both cellular senescence *in vitro* and aging *in vivo* suggests that quiescence deepening likely shares a molecular mechanistic basis with senescence and aging. Given their different degrees of reduced proliferative capacity, deep quiescence may act as a transitional state on the trajectory towards cellular senescence and aging.

## Discussion

Previous studies have shown that the lysosome-autophagy pathway preserves the proliferative capacity of adult stem cells by preventing irreversibly arrested states such as senescence, apoptosis, and terminal differentiation^6, 17, 18, 20^. The present study shows that instead of a simple ON (reversible)—OFF (irreversible) switch, lysosomal function acts as a dimmer switch to continuously modulate quiescence depth (Fig. 5g), playing a previously underappreciated role in adjusting the proliferative capacity of quiescent cells.

Despite increased lysosomal gene expression and biogenesis (Fig. 2e, h), deep than shallow quiescent REF cells exhibited reduced autophagy flux (Fig. 3d). The molecular mechanism responsible for this reduced autophagy in deep quiescent cells remains unclear, but is likely related to similar autophagy decline observed in many types of aged cells^6, 38-40^ and senescent cells^19, 41^. It also remains unclear why lysosomal gene expression and biogenesis in deep quiescent cells are increased to a level that partially but not completely compensate for the reduced autophagy flux. Further increasing lysosomal gene expression and biogenesis to fully compensate the reduced autophagy may become unattainable or too costly for the cell. Alternatively, the “net” lysosomal function after partial compensation may serve as certain adaptive mechanism—e.g., a counter for the serum-starvation duration (i.e., environmental growth restriction), which determines accordingly how cautious the cell will be before committing to exiting quiescence.

Our results suggest that lysosomal function prevents quiescence deepening via ROS reduction (Fig. 5). Previous studies have shown that the lysosome-autophagy pathway reduces ROS in several types of quiescent cells^6, 17, 18, 20^ through mitophagy, i.e., selective autophagic degradation of damaged mitochondria^6^. It remains to be tested whether lysosome function prevents quiescence deepening in REF cells via mitophagy, but the reduced mitochondrial ROS level resulted from enhanced lysosomal function (Fig. 5e, f) is consistent with this model.

It was shown previously that the mammalian target of rapamycin (mTOR) can transform muscle stem cells to a shallower quiescent state called G_Alert_, in which cells are sensitized to growth stimulation^15^. Consistently, continuous mTOR activation led to depletion of certain types of quiescent stem cells *in vivo* by forcing cell cycle reentry^42, 43^. It is well established that mTOR drives cell growth and inhibits the lysosome-autophagy pathway^35, 44^. Accordingly, the effect of mTOR on quiescence depth is likely two-sided: by promoting cell growth, mTOR facilitates quiescence exit and thus reduces quiescence depth; by inhibiting lysosomal function, mTOR drives cells deeper into quiescence. In the G_Alert_ case, where mTOR is activated both before and during stimulation, the net result appears to be shallower quiescence. If we inhibit mTOR activity in quiescent cells before but not during serum stimulation, we would expect an increased lysosomal function (before stimulation) but otherwise largely unaffected cell growth (during stimulation), and thus a net result of shallower quiescence. Indeed, we observed that mTOR inhibition in serum-starved quiescent cells by an inhibitor Torin 1 before (but not during) serum stimulation led to progressively shallower quiescence in Torin 1 treatment duration-dependent manner (Fig. S6c).

The lysosome-autophagy pathway plays an important role in cancer physiology and dormancy^7, 32, 33, 45-47^. Activities of MiT/TFE family members including Tfeb and Mitf are upregulated by overexpression or nuclear localization in multiple cancer types including lung, pancreatic, and ovarian cancers^33, 48, 49^, which is linked to poor prognosis and survival^48, 49^. The lysosome-autophagy pathway may maintain dormant cancer cells at certain quiescence depth that facilitates survival and metastasis^7^. Consistently, autophagy inhibition decreases the viability of dormant breast cancer cells and their metastatic recurrence, suggesting a promising treatment strategy^46^. Future studies are needed to determine the optimal target and degree of lysosomal-autophagy inhibition in treatment to minimize disrupting the quiescence depth of normal cells.

Our study highlights that deep quiescent cells may experience similar gene expression changes as senescent and aged cells (Fig. 6b-e). A shared underlying mechanism may be DNA damage caused by ROS accumulation that is common in these cells. DNA damage is associated with cellular senescence *in vitro*^50-52^ and aging *in vivo*^53, 54^ that are caused by various agents (e.g., ROS and replication stress)^55-57^. DNA damage can induce polyadenylation of replication-dependent histone mRNAs^58^, which are otherwise usually non-polyadenylated^59^. Likely not coincidentally, in both deep quiescent cells and senescent cells, an increase in polyadenylated mRNAs was associated with replication-dependent histones but not replication-independent histones (Fig. S7). In addition to the decrease of lysosomal function and accumulation of ROS and DNA damage, many other cellular activities are up- or down-regulated as cells move deeper in quiescence (Fig. S2c). Some of these cellular activities may also be involved in the regulation of quiescence depth, with their detailed mechanisms awaiting further studies. In this regard, it has been shown recently in both NSCs and bacteria that the accumulation of protein aggregates is associated with quiescence deepening, and that the clearance of protein aggregates (by lysosome in NSCs and by DnaK-ClpB complex in bacteria) enhances the ability of cells to re-enter the cell cycle^31, 60^.

Lastly, it remains elusive how the lysosomal dimmer switch interacts with other cellular control mechanisms of cell growth and arrest. On one end, shallow than deep quiescent cells are more prone to reenter the cell cycle upon grow stimulation, while cell cycle reentry is known to be controlled by the Rb-E2f bistable switch. It has been shown that components of the Rb-E2f-Cyclin/Cdk gene network play important roles in regulating cellular quiescence^12, 61^. We speculate that during quiescence exit, cells first move progressively into shallow quiescence and at a time point (the restriction point^62, 63^) “flip” into the cell cycle by committing to proliferation; the whole process acts like adjusting a dimmer switch before activating a toggle switch. In this regard, it will be important to figure out whether and how the lysosomal switch crosstalks with the Rb-E2f switch in controlling quiescence depth and exit in future studies. On the other end, deep than shallow quiescent cells are more difficult to reenter the cell cycle upon grow stimulation, while the relationship between reversible deep quiescence and irreversible senescence remains mysterious. Our finding that a gene signature for deep quiescence also predicts senescence (Fig. 6b) suggests that shared molecular mechanisms may underlie both cell fates. Relatedly, it is noticeable that the Rb-E2f-p53 gene network, an overlapping mechanism with the one underlying cell cycle entry, controls the entry into senescence^64-68^. Whether and how the lysosomal switch crosstalks with the Rb-E2f-p53 network in regulating the transition from deep quiescence to senescence and whether such a transition is gradual following a continuum or abrupt as controlled by an ultrasensitive or bistable switch-like mechanism^69, 70^ remain significant unanswered questions (Fig. S1a).

## METHODS

### Cell culture and quiescence induction by serum starvation

Rat embryonic fibroblasts used in this study are from a single-cell clone derived from REF52 cells^71^ and contain a stably integrated human E2F1 promoter-driven destabilized EGFP (E2f-GFP) reporter as previously described (REF/E23 cells^26, 72^), except for in Fig. 5e and f, where REF52 cells without the integrated E2f-GFP reporter were used to avoid its interference with transfected GFP-containing expression vectors. Cells were passaged every 2-3 days and maintained at subconfluency in growth medium: Dulbecco’s Modified Eagle’s Medium (DMEM) supplemented with 10% bovine growth serum (BGS; GE Healthcare, SH30541). To induce quiescence, cells were plated at ∼50% confluence on 6-well plates or 100-mm dishes in growth medium for a day, washed twice with DMEM, and cultured in serum-starvation medium (DMEM containing 0.02% BGS) for the indicated duration (≥ 2 days).

### Quiescence-depth assay with E2f-GFP, EdU, or PI readout

To assess quiescence depth, cells were switched from serum-starvation medium to serum-simulation medium (DMEM containing BGS at a gradient of concentrations as indicated) and harvested at the indicated time points by trypsinization. The cell fraction that re-entered the cell cycle was quantified by assessing the profiles of the E2f-GFP reporter, EdU incorporation, or propidium iodide (PI) DNA staining. For the E2f-GFP readout, harvested cells were fixed with 1% formaldehyde in DPBS. For the EdU assay, 1 μM EdU was included in serum-stimulation medium throughout the experiment; harvested cells were subjected to the Click-iT EdU reaction according to the manufacture’s protocol (Invitrogen, C10418/C10340). For the PI assay, harvested cells were lysed in Nuclear Isolation Medium (0.5% bovine serum albumin, 0.1% NP-40, and 1% RNase A in DPBS) containing 5 µg/ml PI. E2f-GFP, EdU, and PI signal intensities in individual cells (∼10,000 cells per sample) were measured using a BD LSRII or Invitrogen Attune Acoustic Focusing flow cytometer; the acquired data were analyzed using FlowJo software (v. 10.3).

### Lysosomal activity and mitochondrial ROS assays

To assess lysosomal mass, cells in serum-starvation medium were incubated with 50 nM LysoTracker Deep Red (Invitrogen, L12492) for 30 minutes. Subsequently, cells were either trypsinized and processed for flow cytometry, or washed with DMEM, placed back in serum-starvation medium, and observed under a Deltavision Elite Microscope (GE Healthcare). To count LysoTracker foci, images from a Cy5 filter were stacked across the Z-axis and binary processed to define foci. Cellular boundaries were manually determined based on images obtained from both POL and Cy5 filters. The foci number within each cell was determined using the particle analysis function in Fiji^73^. To assess lysosomal proteolytic degradation, cells were incubated with 10 µg/ml DQ-Red BSA (Invitrogen, D12051) for an hour and subsequently incubated with or without lysosomal inhibitor for 5.5 hours. Cells were then stained with 2 μM CellTrace Violet (Invitrogen, C34557) for 20 minutes to stain the cell body, washed twice with DMEM, and placed back in serum-starvation medium for Deltavision imaging. To assess mitochondrial ROS level, serum-starved cells were stained with 3.25 μM MitoSox Red (Invitrogen, M36008) for 20 minutes and subsequently trypsinized and processed for flow cytometry.

### Lysosomal function modulation

To inhibit lysosomal function, cells were treated with CQ (chloroquine; Sigma, C6628) or Baf (bafilomycin A1; LC Laboratories, B-1080) added to serum-starvation medium at the indicated concentrations. To enhance lysosomal function, cells were transfected with a human MITF expression vector pEGFP-N1-MITF-A (Addgene, #38132) or control vector (pd2EGFP-N1 from Clontech, or pCMV-mCherry, a gift from Lingchong You) using Neon electroporation (Invitrogen). Briefly, approximately 10^6^ cells with 10 µg plasmid DNA were electroporated in a 100-µl Neon tip with a 20-ms pulse at 1900 V. Cells were plated in 6-well plates or 100-mm dishes at ∼50% confluence and incubated in growth medium for 30 hours to allow recovery. Cells were further cultured in serum-starvation medium for 4 days before assessing the modulation of lysosomal function and quiescence depth by Mitf expression.

### Autophagy flux assay

Autophagy flux was measured by a LC3-II turnover assay, similar to Ref^30^. Briefly, cells were incubated with or without 40 µM CQ for 3-6 hours, washed once with DPBS, snap frozen in liquid nitrogen, and stored in −80 °C until cell lysis. Frozen cells were lysed on ice by RIPA lysis buffer and processed for immunoblot with anti-LC3B antibody (Sigma, L7543), anti-Tubulin alpha antibody (Thermo Scientific, RB-9281-P0), and secondary antibody (LI-COR, 926-68023). Immunoblots were imaged using a LI-COR Odyssey Scanner and analyzed with Fiji software^73^. The LC3-II Δ between CQ-treated and non-treated samples was quantified to reflect autophagy flux^30^.

### cDNA library preparation, RNA-seq and data preprocessing

Total RNA was isolated with a Quick-RNA kit (Zymo Research, R1050). The quality of the RNA (RQN score ≥ 7.5) was confirmed using the Fragment Analyzer platform (Advanced Analytical Technologies). Libraries were prepared using the NEBNext Poly(A) mRNA Magnetic Isolation Module (NEB, E7490S) and NEBNext Ultra RNA Library Prep Kit for Illumina (NEB, E7530L) according to the manufacturer’s instructions. The final quality-ensured libraries were pooled and sequenced on an Illumina HiSeq 2500 for 100 bp paired-end sequencing. Paired-end cleaned reads were aligned to the rat reference genome rn6 (UCSC) using TopHat (v2.1.1) with default parameters^74^. Transcript annotation and normalization to FPKM were handled using Cufflinks (v2.2.1)^74^. Differentially expressed genes between two time points were identified based on fold difference > 2 in FPKM, after filtering out low (FPKM < 8) or inconsistent expression (fold difference > 2 between replicates).

### Gene expression and pathway enrichment analysis of RNA-seq data

To visualize the sequential transition of transcriptome, the expression matrix of differentially expressed genes was log_2_ transformed and subjected to Principal Component Analysis using the R function “prcomp”, with the result visualized using the R package rgl^75^. For gene expression clustering analysis, FPKM was log_2_ transformed and mean-centered on each gene. K-means clustering was performed by Cluster 3.0^76^ and the optimal cluster number was decided by silhouette width. Hierarchical clustering was performed by Cluster 3.0 using the average linkage method. Clustering results were visualized as heat maps using Java Treeview^77^.

Pathways enriched in K-means clusters were analyzed with the DAVID functional annotation tool^78^. Significantly enriched KEGG pathways (p.adj < 0.05) were determined in the KEGG over-representation test using the R package clusterProfiler^79^. Gene Set Enrichment Analysis (GSEA)^80^ was performed to identify gene sets significantly correlated with quiescence depth, run in the “continuous phenotype” mode using the gene set “c2.all.v6.0.symbols.gmt.geneset” from MSigDB^80^ with the sample label corresponding to serum-starvation days (e.g., 2 for 2-day serum starvation). Genes were ranked by Pearson correlation. Identified significant gene sets (FDR ≤ 0.1) were visualized by NetworkX^81^ and Gephi^82^ in a network to resolve gene sets redundancy; two gene sets with a Jaccard index > 0.5 were connected by an edge, and node size was set to reflect the normalized enrichment score (NES).

### TF-target and lysosomal co-expression network construction

To construct a TF-target network, TF-target interactions were downloaded from RegNetwork^83^ and PAZAR^84^ (mouse interactions were used as rat data were unavailable), based on which differentially expressed genes were connected into a directional graph using the Python package NeworkX^81^ and visualized using Gephi^82^ with the Force Atlas mode. The size and color of a node were determined by its betweenness centrality and expression level, respectively.

To construct a lysosomal co-expression network, differentially expressed genes in the form of a log_2_-transformed expression matrix were clustered into co-expression modules using the blockwiseModules function in the R package WGCNA^85^, with the soft-thresholding power and mergeCutHeight set to 20 and 0.25 respectively. The co-expression module containing the largest number of lysosomal genes up-regulated with quiescence deepening was chosen as the lysosomal co-expression network. Genes in the network were connected based on their co-expression degree (i.e., pairwise correlation) with an adjacency threshold of 0.25. Lysosomal genes and TFs were identified using KEGG^86^ and the DBD transcription factor database^87^, respectively. The network was visualized using Cytoscape^88^.

### Quiescence-depth signature model

To identify a gene expression signature reflecting quiescence depth, linear regression with an elastic net penalty was performed on the time-course RNA-seq data (2- to 16-day serum starvation) using the R package penalized^89^, with the sample label set to indicate serum-starvation days (e.g., 2 for 2-day serum starvation). The optimal tuning parameters for L1 and L2 penalties were determined by maximizing the cross-validated log likelihood across the L1 and L2 combinations (0.01 ≤ L1 ≤ 200, 1 ≤ L2 ≤ 100,000). A gene signature reflecting quiescence depth was identified in the resultant regression model. When applied to analyze a given RNA-seq dataset, this regression model generates a corresponding “quiescence depth score” (QDS).

The regression model was applied to previously published transcriptomes related to quiescence^16^, senescence^21, 22^, and aging^23, 24^. The downloaded RNA-seq datasets, if not previously normalized, were normalized to FPKM, RPKM, or CPM with the R packages limma^90^ and edgeR^91^. Gene symbols from human, rat, and mouse were cross-referenced using NCBI HomoloGene, and expression matrices from different species were merged accordingly. Normalized expression values in the merged matrix were log_2_ transformed and mean centered for each tissue or cell type, and analyzed in the regression model above. The resultant QDS reflects the relative “quiescence depth” of the studied tissue or cell type.

## SUPPLEMENTARY FIGURE LEGENDS

**Supplementary Figure 1.**
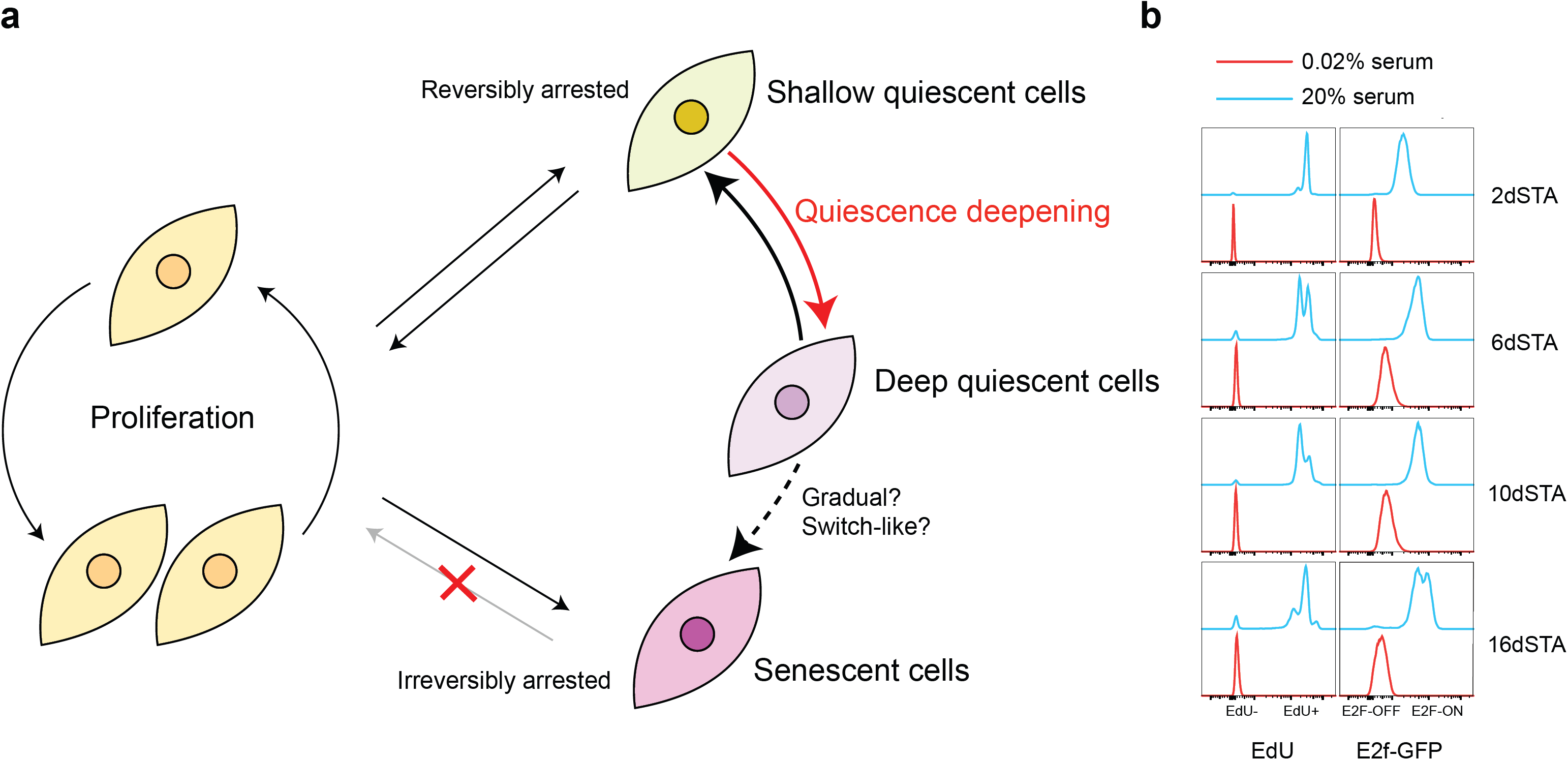
Model of quiescence deepening and deep quiescence as a reversibly arrested cellular state. (**a**) Model of quiescence deepening. The potential sequential transition of cells from shallow to deep quiescence, and eventually into senescence, is discussed later in the paper. (**b**) REF cells were serum starved from 2 to 16 days and then either kept in starvation medium (0.02% serum) or stimulated with 20% serum. Cells were subsequently harvested after 41 hours for E2f-GFP or EdU incorporation profiling (∼10,000 cells per sample, with the highest frequency set to 100% at the y-axis of each histogram).

**Supplementary Figure 2.**
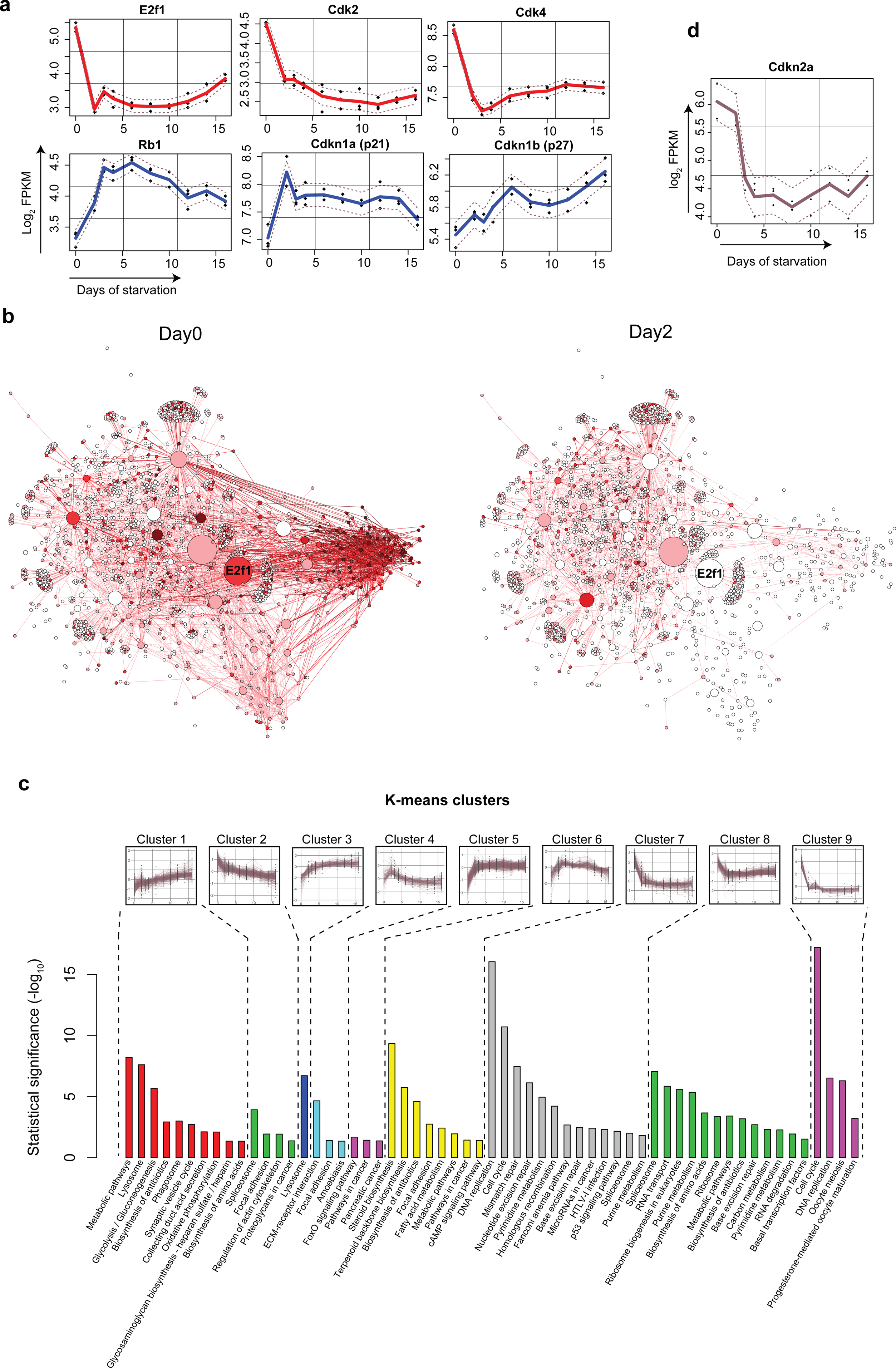
Pro- and anti-proliferative genes were down- and up-regulated respectively in serum-starved cells. (**a**) Time-course expression of E2f1, Cdk2, Cdk4, Rb1, Cdkn1a, and Cdkn1b in RNA-Seq analysis (0-to 16-day serum-starved cells). Dashed line, s.e.m of fitted line. (**b**) Putative transcription factor (TF)-target network of proliferating (day0) and 2-day serum-starved (day2) REF cells. Node color intensity indicates relative gene expression (white to dark red, lowest to highest expression level). Node size indicates the betweenness centrality of the given node in the network. Color of an edge (TF → target) is the same as the color of the target node. (**c**) Pathways significantly enriched in K-means clusters from DAVID functional annotation analysis^78^. Gene expression dynamics in each cluster (same as Fig. 2b) is shown at the top. Y-axis, adjusted p-value calculated by Benjamini correction. (**d**) Time-course expression of Cdkn2a (p16^INK4A^) in RNA-Seq analysis (0-to 16-day serum-starved cells). Dash lines, same as in **a**.

**Supplementary Figure 3.**
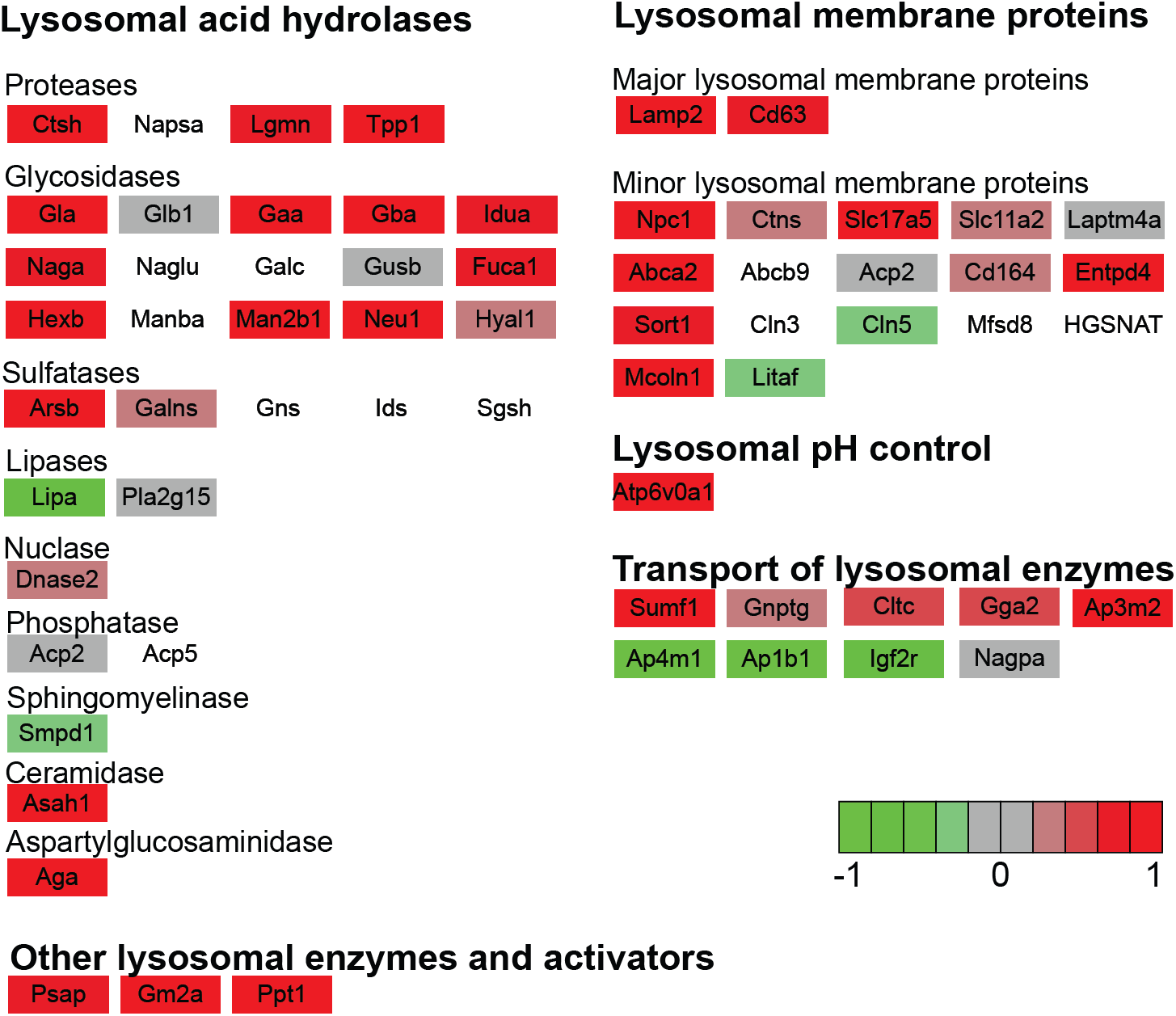
Various categories of lysosomal genes are up-regulated in 16-day serum-starved cells. Up- or down-regulation of various categories of lysosomal genes in 16-day serum-starved cells compared to 2-day serum-starved cells are shown in red or green, respectively, with the degree of changes (log transformed) indicated by the color gradient bar.

**Supplementary Figure 4.**
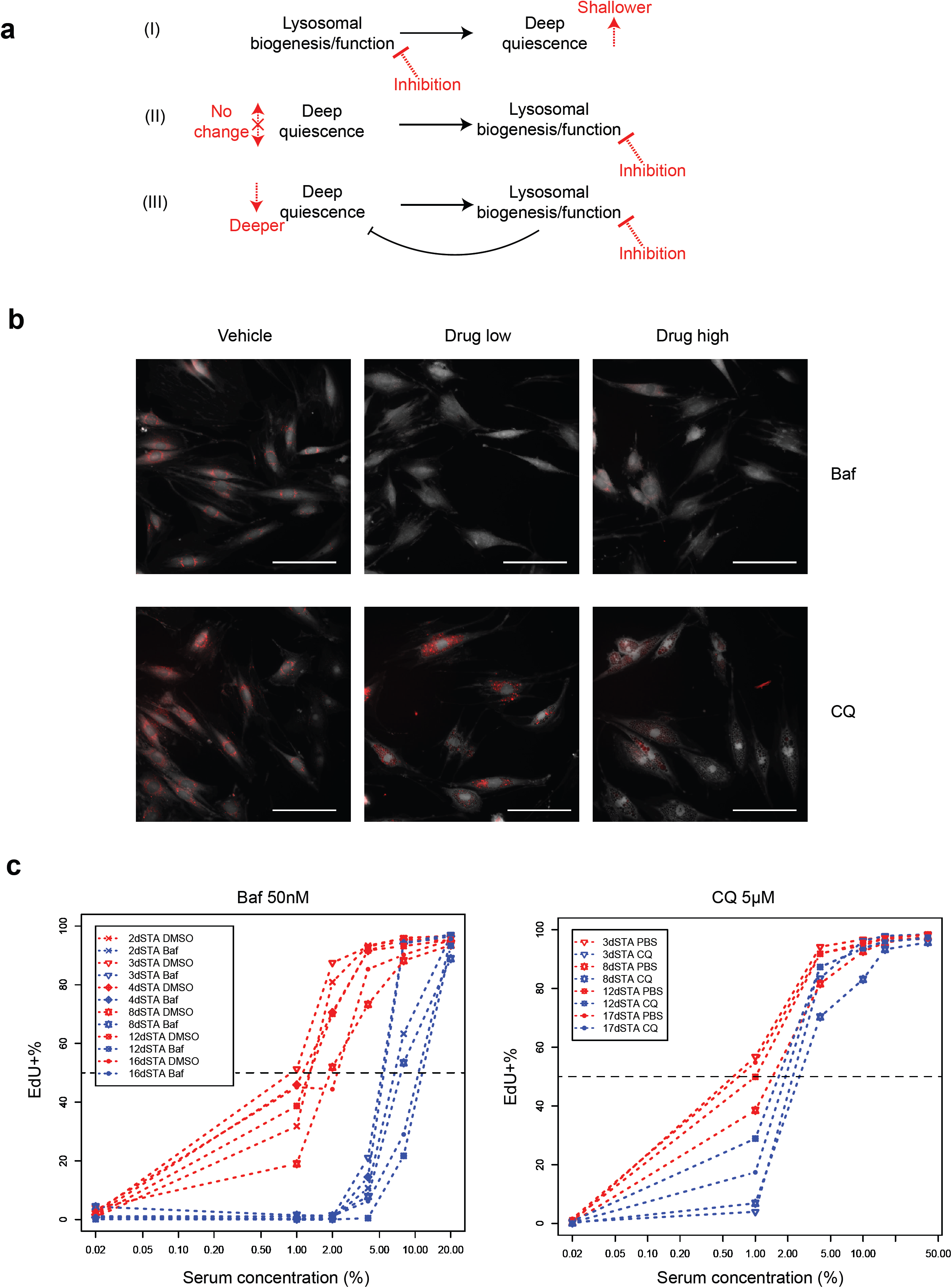
Lysosomal inhibition deepens quiescence depth. (**a**) Three competing hypotheses regarding the relationship between lysosomal biogenesis/function and deep quiescence. See text for details. (**b**) The effects of CQ and Baf on proteolytic degradation. Drug low and Drug high: for CQ, 5 µM and 15 µM, respectively; for Baf, 10 nM and 50 nM, respectively. The degree of proteolytic degradation was indicated by DQ-BSA signal intensity (red puncta); cells were co-stained with CellTrace (gray background stain, see Methods for details). (**c**) Cells serum-starved for different durations were treated with Baf or CQ as in Fig. 3a. Cells were stimulated with serum at indicated concentrations for 40-42 hours and subjected to EdU assay. The EdU+ percentages (y-axis) of drug-treated cells (blue data points) were lower than those of corresponding non-treated controls (red) regardless of serum-starvation days before serum stimulation (see legend box) and serum concentrations applied (> 0.02%, x-axis).

**Supplementary Figure 5.**
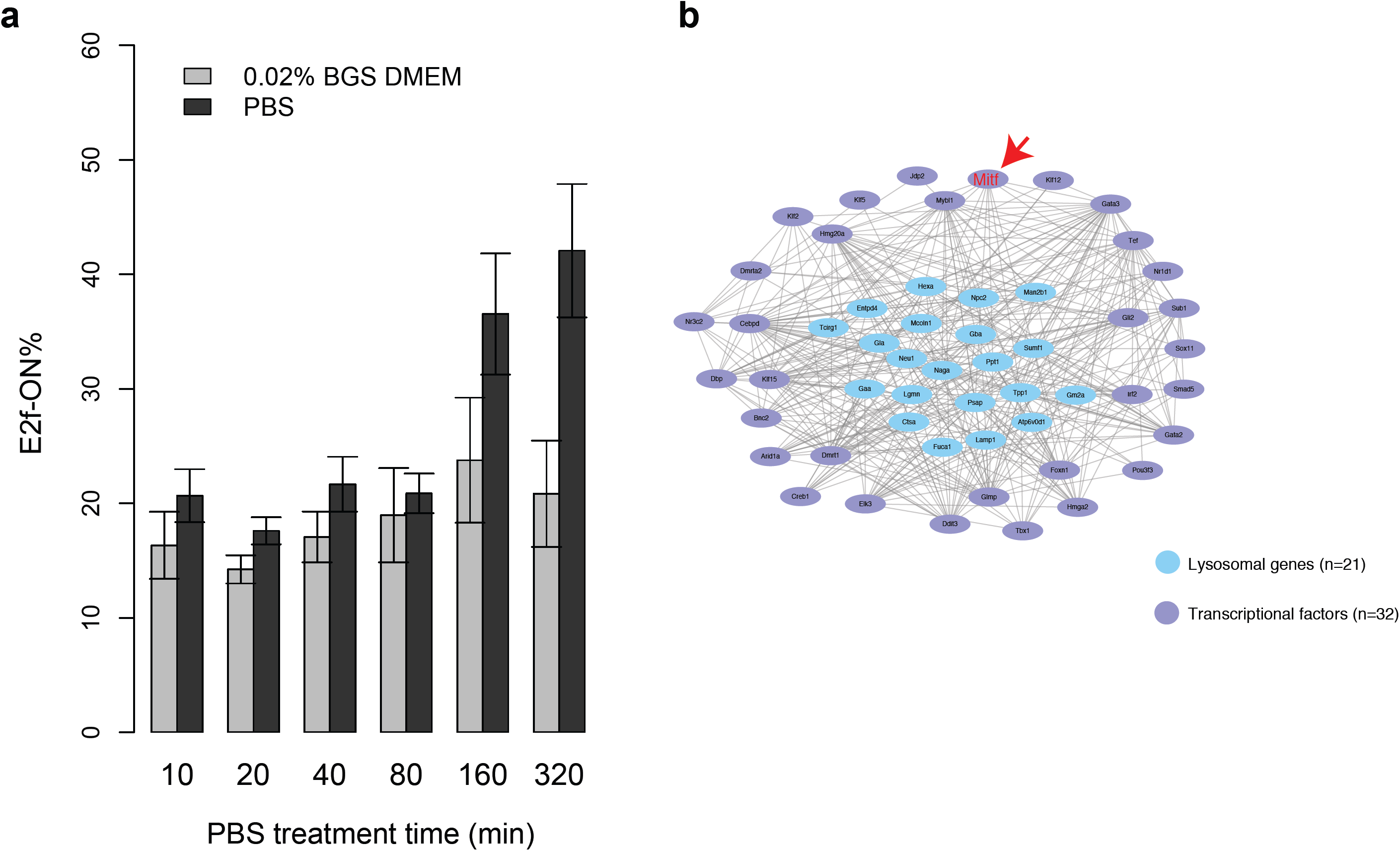
Enhancing lysosomal function pushes cells toward shallower quiescence. (**a**) REF cells were serum starved for 4 days and further cultured in starvation medium (DMEM containing 0.02% BGS) or DPBS for the indicated duration (x-axis). Cells were stimulated with 1% serum in DMEM for 24 hours and measured for E2f-GFP activity (y-axis). Error bar, s.e.m. (**b**) Lysosomal co-expression network associated with deep quiescence. Highly co-expressed lysosomal genes (blue) and TFs (purple) are connected based on the degree of co-expression. See Methods for detail. Mitf is highlighted by a red arrow.

**Supplementary Figure 6.**
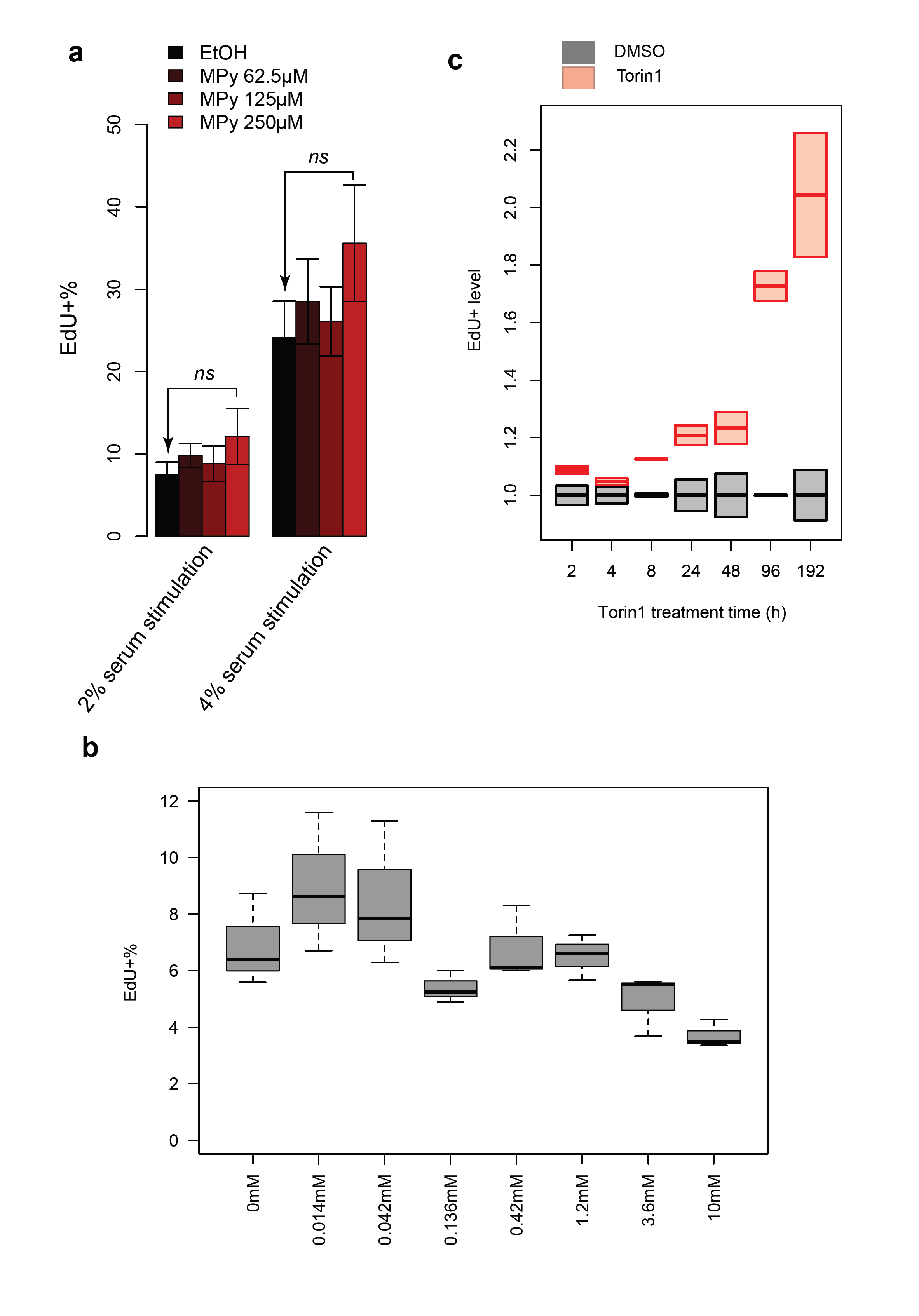
Effects of MPy supplement and mTOR inhibition on quiescence depth. (**a, b**) Effects of MPy supplement. 2-day serum-starved cells were further serum starved with daily supplementation of MPy at indicated concentrations (0-0.25 mM, **a**; 0-10 mM, **b**) for 4 days (**a**) or 2 days (**b**). Cells were stimulated with 2% or 4% serum (**a**) or 1% serum (**b**) for 24 hours and subjected to EdU assay (triplicates). Error bar in **a**, s.e.m. Box plot in **b**, same as in Fig. 2h. (**c**) Effects of mTOR inhibition. 2-day serum-starved cells were further serum starved with daily supplementation of 10 nM Torin1 for the indicated durations. Cells were stimulated with 4% serum for 24 hours and subjected to EdU assay (duplicates). EdU^+^ level (y-axis) corresponds to EdU^+^% in Torin1-treated over non-treated control cells (set to 1.0) in each treatment condition. Bottom and top edges of each box indicate the 1^st^ and 3^rd^ quartiles, respectively.

**Supplementary Figure 7.**
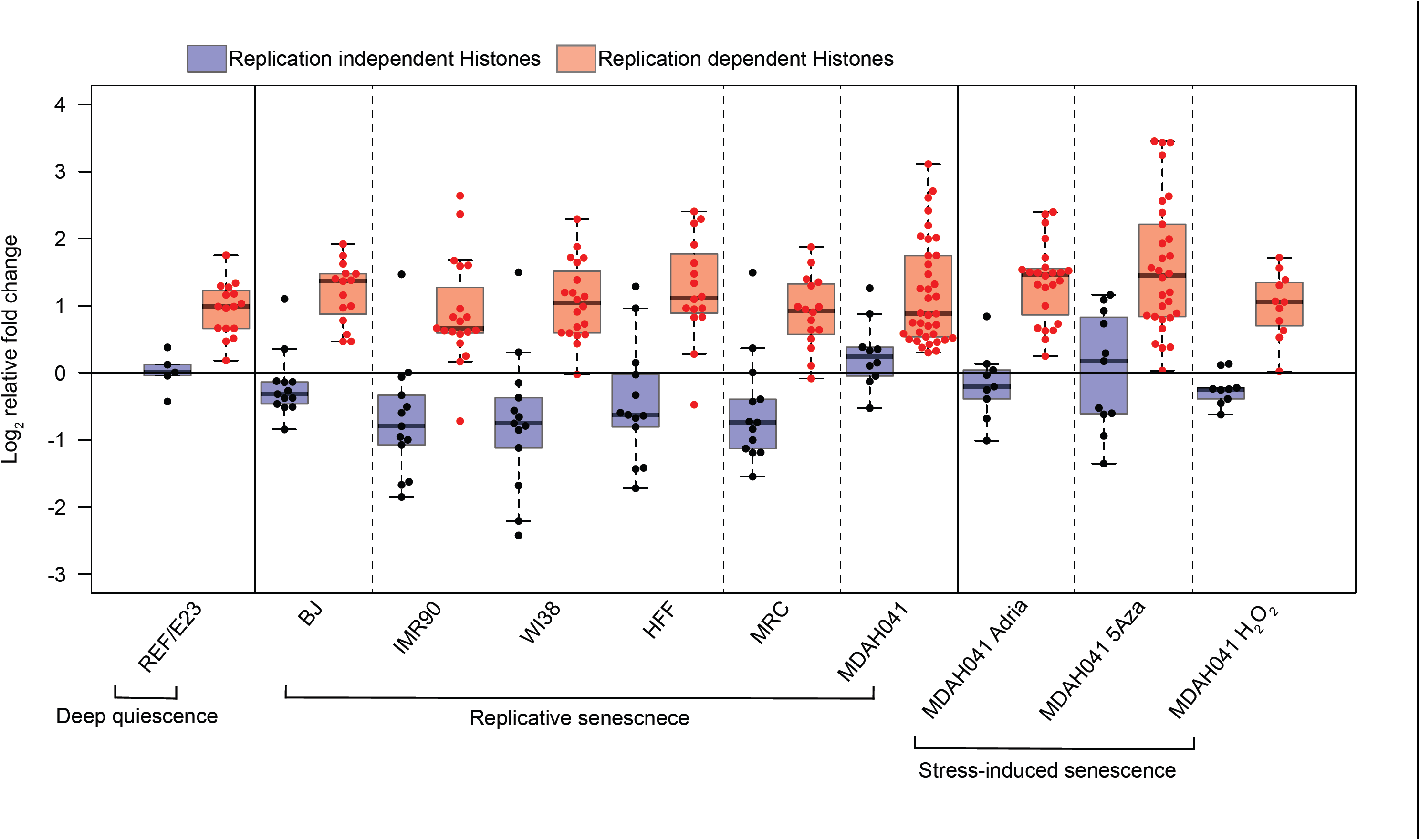
Quiescence deepening are related to DNA damage. Histone genes were identified using the HUGO Gene Nomenclature, and replication-dependent histones were identified according to Ref. 92. Shown are relative log_2_ fold changes of polyadenylated histone mRNAs (replication-dependent and replication-independent) in deep quiescent cells (16-day serum-starved cells) and senescent cells (at indicated conditions) compared to corresponding controls (2-day serum-starved cells and proliferating cells, respectively).

## References

1. Rumman, M., Dhawan, J. & Kassem, M. Concise Review: Quiescence in Adult Stem Cells: Biological Significance and Relevance to Tissue Regeneration. Stem Cells 33, 2903–2912 (2015).

2. Cheung, T.H. & Rando, T.A. Molecular regulation of stem cell quiescence. Nature Reviews Molecular Cell Biology 14, 329 (2013).

3. Rossi, L. et al. Less is more: unveiling the functional core of hematopoietic stem cells through knockout mice. Cell Stem Cell 11, 302–317 (2012).

4. Coller, H.A., Sang, L. & Roberts, J.M. A new description of cellular quiescence. PLoS Biol 4, e83 (2006).

5. Sousa-Victor, P., Garcia-Prat, L., Serrano, A.L., Perdiguero, E. & Munoz-Canoves, P. Muscle stem cell aging: regulation and rejuvenation. Trends Endocrinol Metab 26, 287– 296 (2015).

6. Garcia-Prat, L. et al. Autophagy maintains stemness by preventing senescence. Nature 529, 37–42 (2016).

7. Sosa, M.S., Bragado, P. & Aguirre-Ghiso, J.A. Mechanisms of disseminated cancer cell dormancy: an awakening field. Nat Rev Cancer 14, 611–622 (2014).

8. Oh, J., Lee, Y.D. & Wagers, A.J. Stem cell aging: mechanisms, regulators and therapeutic opportunities. Nat Med 20, 870–880 (2014).

9. Sharpless, N.E. & DePinho, R.A. How stem cells age and why this makes us grow old. Nature Reviews Molecular Cell Biology 8, 703 (2007).

10. Owen, T.A., Soprano, D.R. & Soprano, K.J. Analysis of the growth factor requirements for stimulation of WI-38 cells after extended periods of density-dependent growth arrest. J Cell Physiol 139, 424–431 (1989).

11. Augenlicht, L.H. & Baserga, R. Changes in the G0 state of WI-38 fibroblasts at different times after confluence. Exp Cell Res 89, 255–262 (1974).

12. Kwon, J.S. et al. Controlling Depth of Cellular Quiescence by an Rb-E2F Network Switch. Cell Rep 20, 3223–3235 (2017).

13. Bucher, N.L. Regeneration of Mammalian Liver. Int Rev Cytol 15, 245–300 (1963).

14. Roth, G.S. & Adelman, R.C. Age-dependent regulation of mammalian DNA synthesis and cell division in vivo by glucocorticoids. Exp Gerontol 9, 27–31 (1974).

15. Rodgers, J.T. et al. mTORC1 controls the adaptive transition of quiescent stem cells from G0 to G(Alert). Nature 510, 393–396 (2014).

16. Llorens-Bobadilla, E. et al. Single-Cell Transcriptomics Reveals a Population of Dormant Neural Stem Cells that Become Activated upon Brain Injury. Cell Stem Cell 17, 329–340 (2015).

17. Warr, M.R. et al. FOXO3A directs a protective autophagy program in haematopoietic stem cells. Nature 494, 323–327 (2013).

18. Ho, T.T. et al. Autophagy maintains the metabolism and function of young and old stem cells. Nature 543, 205–210 (2017).

19. Kang, H.T., Lee, K.B., Kim, S.Y., Choi, H.R. & Park, S.C. Autophagy impairment induces premature senescence in primary human fibroblasts. PLoS One 6, e23367 (2011).

20. Lum, J.J. et al. Growth factor regulation of autophagy and cell survival in the absence of apoptosis. Cell 120, 237–248 (2005).

21. Purcell, M., Kruger, A. & Tainsky, M.A. Gene expression profiling of replicative and induced senescence. Cell Cycle 13, 3927–3937 (2014).

22. Marthandan, S. et al. Conserved Senescence Associated Genes and Pathways in Primary Human Fibroblasts Detected by RNA-Seq. PLoS One 11, e0154531 (2016).

23. Grover, A. et al. Single-cell RNA sequencing reveals molecular and functional platelet bias of aged haematopoietic stem cells. Nat Commun 7, 11075 (2016).

24. Yu, Y. et al. A rat RNA-Seq transcriptomic BodyMap across 11 organs and 4 developmental stages. Nat Commun 5, 3230 (2014).

25. Yao, G., Tan, C., West, M., Nevins, J.R. & You, L. Origin of bistability underlying mammalian cell cycle entry. Mol Syst Biol 7, 485 (2011).

26. Yao, G., Lee, T.J., Mori, S., Nevins, J.R. & You, L. A bistable Rb-E2F switch underlies the restriction point. Nat Cell Biol 10, 476–482 (2008).

27. Sharpless, N.E. & Sherr, C.J. Forging a signature of in vivo senescence. Nat Rev Cancer 15, 397–408 (2015).

28. Nazio, F. et al. Fine-tuning of ULK1 mRNA and protein levels is required for autophagy oscillation. J Cell Biol 215, 841–856 (2016).

29. Klionsky, D.J. et al. Guidelines for the use and interpretation of assays for monitoring autophagy (3rd edition). Autophagy 12, 1–222 (2016).

30. Mizushima, N., Yoshimori, T. & Levine, B. Methods in mammalian autophagy research. Cell 140, 313–326 (2010).

31. Leeman, D.S. et al. Lysosome activation clears aggregates and enhances quiescent neural stem cell activation during aging. Science 359, 1277–1283 (2018).

32. Ploper, D. et al. MITF drives endolysosomal biogenesis and potentiates Wnt signaling in melanoma cells. Proc Natl Acad Sci U S A 112, E420–429 (2015).

33. Perera, R.M. et al. Transcriptional control of autophagy-lysosome function drives pancreatic cancer metabolism. Nature 524, 361–365 (2015).

34. Bouche, V. et al. Drosophila Mitf regulates the V-ATPase and the lysosomal-autophagic pathway. Autophagy 12, 484–498 (2016).

35. Settembre, C., Fraldi, A., Medina, D.L. & Ballabio, A. Signals from the lysosome: a control centre for cellular clearance and energy metabolism. Nat Rev Mol Cell Biol 14, 283–296 (2013).

36. Rubinsztein, David C., Mariño, G. & Kroemer, G. Autophagy and Aging. Cell 146, 682– 695 (2011).

37. Hernandez-Segura, A. et al. Unmasking Transcriptional Heterogeneity in Senescent Cells. Curr Biol 27, 2652–2660 e2654 (2017).

38. Vittorini, S. et al. The age-related accumulation of protein carbonyl in rat liver correlates with the age-related decline in liver proteolytic activities. J Gerontol A Biol Sci Med Sci 54, B318–323 (1999).

39. Kaushik, S. et al. Loss of autophagy in hypothalamic POMC neurons impairs lipolysis. EMBO Rep 13, 258–265 (2012).

40. Cuervo, A.M. & Dice, J.F. Age-related decline in chaperone-mediated autophagy. J Biol Chem 275, 31505–31513 (2000).

41. Han, B.I., Hwang, S.H. & Lee, M. A progressive reduction in autophagic capacity contributes to induction of replicative senescence in Hs68 cells. Int J Biochem Cell Biol 92, 18–25 (2017).

42. Chen, C. et al. TSC-mTOR maintains quiescence and function of hematopoietic stem cells by repressing mitochondrial biogenesis and reactive oxygen species. J Exp Med 205, 2397–2408 (2008).

43. Adhikari, D. et al. Tsc/mTORC1 signaling in oocytes governs the quiescence and activation of primordial follicles. Hum Mol Genet 19, 397–410 (2010).

44. Laplante, M. & Sabatini, D.M. mTOR signaling in growth control and disease. Cell 149, 274–293 (2012).

45. Yang, A. & Kimmelman, A.C. Inhibition of autophagy attenuates pancreatic cancer growth independent of TP53/TRP53 status. Autophagy 10, 1683–1684 (2014).

46. Vera-Ramirez, L., Vodnala, S.K., Nini, R., Hunter, K.W. & Green, J.E. Autophagy promotes the survival of dormant breast cancer cells and metastatic tumour recurrence. Nat Commun 9, 1944 (2018).

47. Mowers, E.E., Sharifi, M.N. & Macleod, K.F. Autophagy in cancer metastasis. Oncogene 36, 1619–1630 (2017).

48. Giatromanolaki, A. et al. Increased expression of transcription factor EB (TFEB) is associated with autophagy, migratory phenotype and poor prognosis in non-small cell lung cancer. Lung Cancer 90, 98–105 (2015).

49. Fu, X., Zhang, L., Dan, L., Wang, K. & Xu, Y. Expression of TFEB in epithelial ovarian cancer and its role in autophagy. International Journal of Clinical and Experimental Pathology 9, 10914–10928 (2016).

50. Gire, V., Roux, P., Wynford-Thomas, D., Brondello, J.M. & Dulic, V. DNA damage checkpoint kinase Chk2 triggers replicative senescence. EMBO J 23, 2554–2563 (2004).

51. Zhang, H. Molecular signaling and genetic pathways of senescence: Its role in tumorigenesis and aging. J Cell Physiol 210, 567–574 (2007).

52. d’Adda di Fagagna, F. Living on a break: cellular senescence as a DNA-damage response. Nature Reviews Cancer 8, 512 (2008).

53. Soares, J.P. et al. Aging and DNA damage in humans: a meta-analysis study. Aging (Albany NY) 6, 432–439 (2014).

54. Beerman, I., Seita, J., Inlay, Matthew A., Weissman, Irving L. & Rossi, Derrick J. Quiescent Hematopoietic Stem Cells Accumulate DNA Damage during Aging that Is Repaired upon Entry into Cell Cycle. Cell Stem Cell 15, 37–50 (2014).

55. Cooke, M.S., Evans, M.D., Dizdaroglu, M. & Lunec, J. Oxidative DNA damage: mechanisms, mutation, and disease. The FASEB Journal 17, 1195–1214 (2003).

56. Venkatachalam, G., Surana, U. & Clement, M.V. Replication stress-induced endogenous DNA damage drives cellular senescence induced by a sub-lethal oxidative stress. Nucleic Acids Res 45, 10564–10582 (2017).

57. Techer, H., Koundrioukoff, S., Nicolas, A. & Debatisse, M. The impact of replication stress on replication dynamics and DNA damage in vertebrate cells. Nat Rev Genet 18, 535–550 (2017).

58. Kari, V. et al. A subset of histone H2B genes produces polyadenylated mRNAs under a variety of cellular conditions. PLoS One 8, e63745 (2013).

59. Marzluff, W.F., Wagner, E.J. & Duronio, R.J. Metabolism and regulation of canonical histone mRNAs: life without a poly(A) tail. Nature reviews. Genetics 9, 843–854 (2008).

60. Pu, Y. et al. Dynamic protein aggregation regulates bacterial dormancy depth critical for antibiotic tolerance. bioRxiv (2017).

61. Spencer, Sabrina L. et al. The Proliferation-Quiescence Decision Is Controlled by a Bifurcation in CDK2 Activity at Mitotic Exit. Cell 155, 369–383 (2013).

62. Schwarz, C. et al. A Precise Cdk Activity Threshold Determines Passage through the Restriction Point. Molecular Cell 69, 253–264.e255 (2018).

63. Matson, J.P. & Cook, J.G. Cell cycle proliferation decisions: the impact of single cell analyses. FEBS J 284, 362–375 (2017).

64. Childs, B.G., Durik, M., Baker, D.J. & van Deursen, J.M. Cellular senescence in aging and age-related disease: from mechanisms to therapy. Nature Medicine 21, 1424 (2015).

65. Sherr, C.J. & DePinho, R.A. Cellular Senescence. Cell 102, 407–410 (2000).

66. Shay, J.W., Pereira-Smith, O.M. & Wright, W.E. A role for both RB and p53 in the regulation of human cellular senescence. Experimental Cell Research 196, 33–39 (1991).

67. Campisi, J. & d’Adda di Fagagna, F. Cellular senescence: when bad things happen to good cells. Nature Reviews Molecular Cell Biology 8, 729 (2007).

68. Bennett, D.C. Human melanocyte senescence and melanoma susceptibility genes. Oncogene 22, 3063 (2003).

69. Tyson, J.J., Chen, K.C. & Novak, B. Sniffers, buzzers, toggles and blinkers: dynamics of regulatory and signaling pathways in the cell. Current Opinion in Cell Biology 15, 221– 231 (2003).

70. Ferrell, J.E. & Ha, S.H. Ultrasensitivity part III: cascades, bistable switches, and oscillators. Trends in Biochemical Sciences 39, 612–618 (2014).

71. Logan, J. et al. Transformation by adenovirus early region 2A temperature-sensitive mutants and their revertants. Virology 115, 419–422 (1981).

72. Wang, X. et al. Exit from quiescence displays a memory of cell growth and division. Nat Commun 8, 321 (2017).

73. Schindelin, J. et al. Fiji: an open-source platform for biological-image analysis. Nat Methods 9, 676–682 (2012).

74. Trapnell, C. et al. Differential gene and transcript expression analysis of RNA-seq experiments with TopHat and Cufflinks. Nat Protoc 7, 562–578 (2012).

75. Adler, D., Murdoch, D. & others R package version 0.99.16. (2018).

76. de Hoon, M.J.L., Imoto, S., Nolan, J. & Miyano, S. Open source clustering software. Bioinformatics 20, 1453–1454 (2004).

77. Saldanha, A.J. Java Treeview-extensible visualization of microarray data. Bioinformatics 20, 3246–3248 (2004).

78. Huang, D.W., Sherman, B.T. & Lempicki, R.A. Systematic and integrative analysis of large gene lists using DAVID bioinformatics resources. Nature Protocols 4, 44 (2008).

79. Yu, G., Wang, L.-G., Han, Y. & He, Q.-Y. clusterProfiler: an R Package for Comparing Biological Themes Among Gene Clusters. OMICS: A Journal of Integrative Biology 16, 284–287 (2012).

80. Subramanian, A. et al. Gene set enrichment analysis: a knowledge-based approach for interpreting genome-wide expression profiles. Proc Natl Acad Sci U S A 102, 15545– 15550 (2005).

81. Hagberg, A., Swart, P. & S Chult, D. in Proceedings of the 7th Python in Science Conference (SciPy 2008) (2008).

82. Bastian, M., Heymann, S. & Jacomy, M. in International AAAI Conference on Weblogs and Social Media (2009).

83. Liu, Z.P., Wu, C., Miao, H. & Wu, H. RegNetwork: an integrated database of transcriptional and post-transcriptional regulatory networks in human and mouse. Database (Oxford) 2015 (2015).

84. Portales-Casamar, E. et al. PAZAR: a framework for collection and dissemination of cis-regulatory sequence annotation. Genome Biol 8, R207 (2007).

85. Langfelder, P. & Horvath, S. WGCNA: an R package for weighted correlation network analysis. BMC Bioinformatics 9, 559 (2008).

86. Kanehisa, M., Furumichi, M., Tanabe, M., Sato, Y. & Morishima, K. KEGG: new perspectives on genomes, pathways, diseases and drugs. Nucleic Acids Research 45, D353–D361 (2017).

87. Kummerfeld, S.K. & Teichmann, S.A. DBD: a transcription factor prediction database. Nucleic Acids Res 34, D74–81 (2006).

88. Shannon, P. et al. Cytoscape: a software environment for integrated models of biomolecular interaction networks. Genome Res 13, 2498–2504 (2003).

89. Goeman, J.J. L1 penalized estimation in the Cox proportional hazards model. Biom J 52, 70–84 (2010).

90. Ritchie, M.E. et al. limma powers differential expression analyses for RNA-sequencing and microarray studies. Nucleic Acids Res 43, e47 (2015).

91. Robinson, M.D., McCarthy, D.J. & Smyth, G.K. edgeR: a Bioconductor package for differential expression analysis of digital gene expression data. Bioinformatics 26, 139– 140 (2010).

92. Marzluff, W.F., Gongidi, P., Woods, K.R., Jin, J. & Maltais, L.J. The human and mouse replication-dependent histone genes. Genomics 80, 487–498 (2002).

